# Improved Efficiency of Two-Step Amplicon PCR Using an Acoustic Liquid Handler

**DOI:** 10.1101/2024.12.06.627172

**Authors:** Brooke R. Benz, Eglantina Lopez Echartea, Briana K. Whitaker, Thomas Baldwin, Barney A. Geddes

## Abstract

2.

The improvement in next-generation sequencing technologies has reduced the costs of sequencing significantly. However, library preparation costs for amplicon sequencing have remained largely unchanged – which is ultimately the cost-limiting step in processing large numbers of microbiome samples. Acoustic liquid handlers can transfer volumes as low as 2.5 nL and have been used to miniaturize several different molecular and cellular assays, including single-step polymerase chain reaction (PCR) amplicon library preparations. However, there are no current methods available for a two-step library preparation process using an acoustic liquid handler. In this study, we tested the efficiency of an acoustic liquid handler to automate the PCRs and library quantification while also incorporating automated library bead cleanup. We compared the material usage and costs for library preparation and sequencing results of this automated method to the standard, manual method. The automated protocol was able to reduce both PCR reaction volumes five-fold and increased efficiency for library preparation by ∼32% without affecting bacterial community compositions. The associated increase in efficiency of our automated method will allow for greater throughput in sequencing hundreds of microbiome samples without affecting the quality of those sequences.

**Impact statement:** Library preparation for amplicon sequencing in microbiome studies remains a significant cost constraint, even while sequencing costs have decreased in the last few decades. Acoustic liquid handling robots have been utilized to miniaturize several different assays, including a one-step PCR library preparation process, to reduce costs. However, there are currently no methods available for utilizing acoustic liquid handlers for two-step PCR library preparation. Here, we show that the incorporation of an acoustic liquid handler and automated purification instruments into a two-step library preparation protocol led to a reduction in costs and time to prepare libraries. This method is comparable to the standard manual method and will lead to significant cost savings for the preparation of hundreds of samples required for microbiome studies.

**Data summary:** All protocols, sequence data, and analysis codes have been made publicly available in open-access repositories. The authors confirm all supporting data, code, and protocols have been provided within the article or through supplementary data files.

Sequencing data was deposited in the NCBI SRA under the BioProject PRJNA1190462.

R scripts used for data analysis are available at https://github.com/NDSU-Geddes-Lab/brb-libprep doi: 10.5281/zenodo.14194699.

Protocols for automated library preparation have been published on protocols.io and are available at dx.doi.org/10.17504/protocols.io.e6nvwb5r2vmk/v2 and dx.doi.org/10.17504/protocols.io.e6nvwb5w2vmk/v2.

## 5. Introduction

There has been a significant increase of microbiome studies across many systems due to the development of Next Generation Sequencing (NGS) technologies. These studies aim to characterize the microbiomes of humans,^1^ plants,^2,3^ animals,^4,5^ and the environment^6^ ranging from soil^7,8^ to marine^9^ ecosystems. NGS has made it possible to sequence hundreds to thousands of samples. Since the development of NGS technologies, there has been a significant reduction in sequencing costs, improvement in sequencing quality, and increased sample read depth per run.^10^ While sequencing capacity and accessibility have greatly improved, there is still a significant bottleneck associated with user-time as well as materials and expenses for the library preparation of these samples.^11^ Even with an overall reduction in sequencing expenses, performing microbiome studies still requires a substantial investment, with prices for library preparation ranging from ∼$30 to $80 per sample.^12,13^ To efficiently process these samples, minimizing reaction volumes, reducing reagent usage and preparation time, and utilizing high-throughput methods are needed.

Hundreds to thousands of samples are required to characterize microbiomes. However, the significant expense to prepare libraries can limit the sample size in these studies. Technological advancements, like the development of automated pipette-based liquid handlers and acoustic liquid handlers that enable the transfer of small volumes, have allowed for the miniaturization of many processes. Acoustic liquid handlers differ from automated pipette-based liquid handlers in that they use sound waves to transfer precise volumes, thus further saving on pipette tip waste. In this study, we evaluated the efficiency and reproducibility of library preparation in nano-volume reactions using an acoustic liquid handler.^14^ Miniaturization of different assays through acoustic liquid handling has successfully been incorporated into a variety of different laboratory processes, including – drug discovery and dosage testing,^15^ mass spectrometry,^16^ plasmid sequencing,^17^ qRT-PCR,^18^ antigen isolation,^19^ and cell-free protein synthesis (CFPS).^20^ Thus, acoustic liquid handlers can be incorporated into a variety of different laboratory processes.

Acoustic liquid handlers were previously used to miniaturize amplicon library preparation as part of the Earth Microbiome Project (EMP) protocol.^21^ In this protocol, a one-step PCR preparation was performed and compared to the standard protocol involving pipette-based liquid handler (i.e., the Eppendorf epMotion). Reducing the reaction volumes, reagents, pipettes, and other consumables significantly decreased the costs of amplicon library preparation. However, the Illumina standard protocol for amplicon library preparation uses a two-step PCR process.^22^ The two-step library preparation method involves two separate PCR steps: an amplicon PCR and an index PCR. The amplicon PCR generates amplicons by targeting specific regions such as the V4 region of the 16S rRNA gene for bacteria or the ITS1 region of the rRNA intergenic region for fungi. Next, the index PCR adds unique barcode sequences and Illumina sequencing adapters to the amplicons. It has been shown that this two-step preparation method is more accurate and reduces primer bias.^23^ Additionally, there has also been growing interest in using a second set of barcodes during the first round of PCR to increase the number of samples in a single sequencing run.^24^ Thus, the two-step PCR approach is likely more amenable to greater throughput than the one-step approach.

In this study, we explored the incorporation of an acoustic liquid handler into the standard two-step library preparation workflows and developed a new protocol for its use. The protocol was applied to a set of microbiome samples from the plant phyllosphere that represent a complex, heterogeneous sample set due to the low microbial loads and the need to block plant organelle DNA with PNAs.^25,26^ We compared our newly developed protocol to the manual preparation and tracked the efficiency of the usage of consumables, reagents, and user-time for each step. We also compared the performance of the automated method when sequenced on an Illumina MiSeq versus Illumina NextSeq to compare sequencing results given the expected read depth disparities. Ultimately, the automated protocol led to a significant reduction in costs and user-time for library preparation compared to the manual protocol.

## 6. Theory and implementation

### Methods

#### Sampling and sample preparation

As a test set for this protocol, we utilized samples collected from a project investigating the impact of *Fusarium* Head Blight (FHB) disease on the barley microbiome. Ten breeding lines were randomly selected from the USDA-ARS breeding lines in the Aberdeen malting barley training population (Supp. Fig. 1A). All ten lines were planted in 2022 at each of three misted FHB nurseries located in Fargo, ND; St. Paul, MN; and Ithaca, NY (Supp. Fig. 1B).

FHB symptoms typically develop 2-4 weeks post-flowering, therefore samples were collected 3-5 weeks after flowering for each location. Five spikes showing visible disease symptoms (labeled “Diseased”) and five spikes showing <30% visible symptoms (labeled “Non-diseased”) were collected for each genotype. Individual spikes were cut from the stem then transported in plastic bags amended with Drierite desiccant granules and kept on ice. Samples were stored at −20°C until further processing.

The spikes were placed in a freeze dryer (Labconco, Kansas City, MO, USA) for 48 to 72 hours. Each spike was then individually ground using a laboratory mill grinder (Perten Instruments, Shelton, CT, USA). To minimize cross-contamination, samples of the same genotype and disease state were ground sequentially. Between each spike, the grinder was cleaned by brushing and vacuuming the leftover ground spike material. Between treatments, the grinder was also cleaned with ethanol. The ground material of each spike was collected in individual 15 mL conical tubes and stored at −20°C.

DNA extractions were performed on 100 milligrams of ground spike material using 96-well DNA extraction kits (Norgen Biotek, Thorold, Ontario, CAN). Each plate contained three controls (DNA extraction negative, PCR negative, and Positive control (ZymoBIOMICS Microbial Community Standard, Irvine, CA, USA)). DNA concentrations were measured with Quant-iT PicoGreen dsDNA Assay Kits (Invitrogen, Waltham, MA, USA) and a Biotek Cytation5 (Agilent Technologies, Santa Clara, CA, USA), then diluted to 5 ng μL^−1^.

#### Bacterial 16S rRNA amplicon library preparation comparison

A random subset of the field samples described above, amounting to one 96-well plate of samples, was prepared using both methods. Figure 1 depicts a comparison of the steps involved in the manual and automated library preparation protocols. Broadly, a two-step PCR approach was performed on the V4 region of the 16S rRNA gene to profile bacterial communities. Following conventional practices,^21^ appropriate checks for successful PCR reactions and amplicon size were made on a random subset of samples for both the manual and automated methods using gel electrophoresis and TapeStation (Agilent Technologies, Santa Clara, CA, USA), respectively.

**Figure 1.**
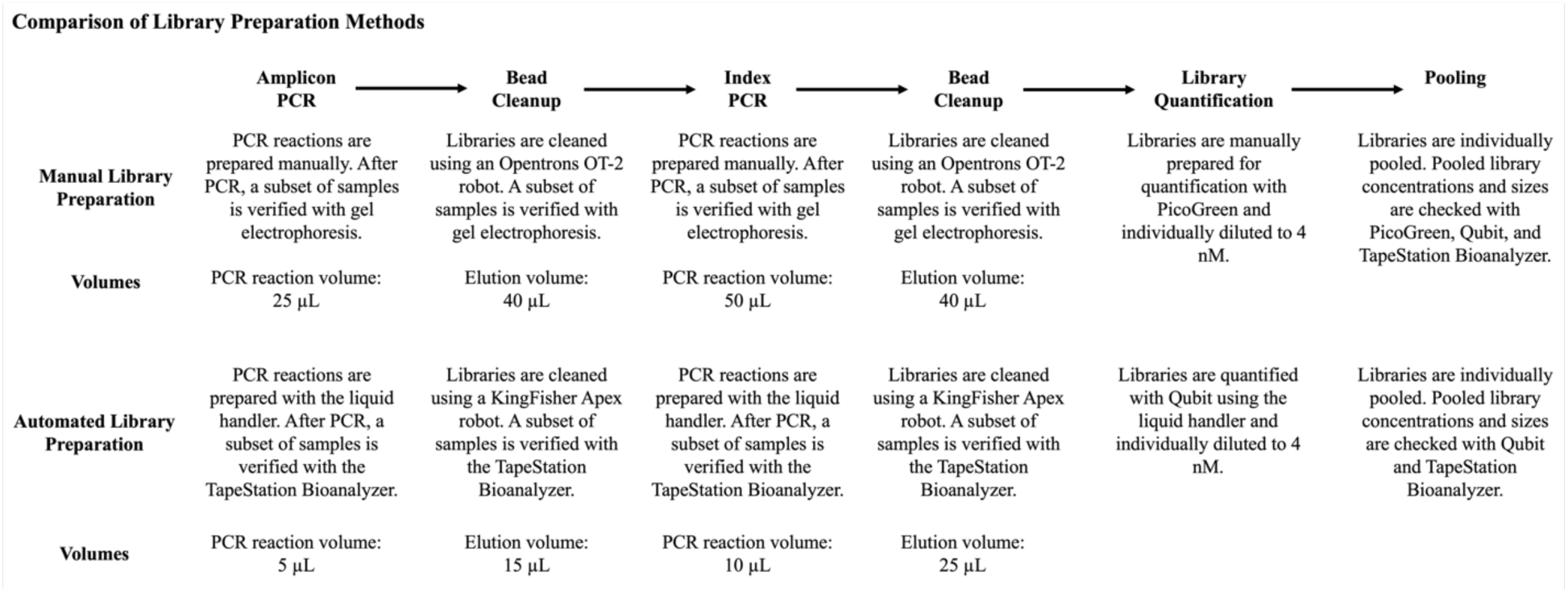
The automated library preparation protocol miniaturized and increased the throughput of library preparation. Comparison of steps between manual and automated library preparation.

The manual amplicon library preparation was performed following a modified version of the Illumina “16S Metagenomic Sequencing Library” protocol.^22^ Individual amplicon PCR reactions consisted of a total reaction volume of 25 μL with 2.5 μL of each forward and reverse V4 amplicon primers (final concentration 0.2 μM; Integrated DNA Technologies Coralville, IA, USA), 12.5 μL of KAPA HiFi DNA polymerase (final concentration 1x; Roche Diagnostics, Indianapolis, IN, USA), 2.5 μL of each mPNA and pPNA blockers (final concentration 2 μM; PNABio, Inc., Thousand Oaks, CA, USA), and 2.5 μL of template DNA. Nextera primer sequences: V4_515F: TCGTCGGCAGCGTCAGATGTGTATAAGAGACAGGTGCCAGCMGCCGCGGTAA; V4_R: GTCTCGTGGGCTCGGAGATGTGTATAAGAGACAGGACTACHVGGGTATCTAATCC.

Amplicon PCR was performed on a thermocycler (Mastercycler epGradient Eppendorf, Enfield, CT, USA) using the following program: 95°C for 3 minutes; 28 cycles of 95°C for 30 seconds, 55°C for 30 seconds, 72°C for 30 seconds; 72°C for 5 minutes; hold at 4°C. Libraries were cleaned with 20 μL of Mag-Bind TotalPure NGS magnetic beads (Omega Bio-tek, Inc., Norcross, GA, USA) using an Opentrons OT-2 robot (Opentrons Labworks, Long Island City, NY, USA). The Opentrons robot was programmed to perform bead cleanup following the procedure in the “16S Metagenomic Sequencing Library” protocol.^22^

Next, index PCR consisted of a total reaction volume of 50 μL and was performed with 5 μL of each forward and reverse index primers (final concentration 0.1 μM; Nextera primer sequences, Integrated DNA Technologies Coralville, IA, USA), 25 μL of DNA polymerase (final concentration 1x), 10 μL of water, and 5 μL of template DNA. Index PCR was performed on a thermocycler using the same program as for amplicon PCR, except 11 cycles instead of 28 cycles. Magnetic bead cleanup was completed after index PCR on the Opentrons robot with the same steps as before except 56 μL of beads were added to each sample. Libraries were quantified using Quant-iT PicoGreen dsDNA Assay Kits (Invitrogen, Waltham, MA, USA), diluted to 4 nM, and pooled together for sequencing.

The automated library preparation method followed the same basic steps in the library preparation process but with a focus on automation and reduced sample volumes. First, the total reaction volume for amplicon PCR was 5 μL. However, all PCR reagents were included at the same final concentrations, including the template DNA. In the automated method, the mastermix and template DNA were transferred to a 384-well PCR plate using the Echo 525 acoustic liquid handler (Beckman Coulter, Indianapolis, IN, USA) instead of pipettes. Amplicon PCR was performed with an Applied Biosystems QuantStudio 6 Flex Real-Time PCR machine (Invitrogen, Waltham, MA, USA) using the following program: 95°C for 3 minutes; 30 cycles of 95°C for 30 seconds, 55°C for 30 seconds, 72°C for 30 seconds; 72°C for 5 minutes; hold at 4°C. An additional two cycles were performed to ensure sufficient amplification with the smaller DNA template volume added. Instead of bead cleanup with the Opentrons robot, individual libraries were cleaned using the KingFisher Apex (Thermo Fisher Scientific, Waltham, MA, USA) as follows. A 96-well sample plate was prepared by combining 10 μL of Mag-Bind TotalPure NGS magnetic beads, 5 μL of molecular-grade water and the libraries. To clean the libraries, two 96-well wash plates were prepared by adding 30 μL of freshly prepared 70% EtOH. To elute the cleaned libraries, 15 μL of molecular-grade water was added to a 96-well elution plate.

Next, the Echo 525 liquid handler was used to transfer the index PCR mastermix and indices to a 384-well PCR plate. Template DNA was manually added to each well. Individual index PCR reactions consisted of 1 μL each of forward and reverse Nextera indices, 5 μL of DNA polymerase, 1 μL of molecular-grade water, and 2 μL of template DNA, for a total reaction volume of 10 μL. Index PCR was performed with an Applied Biosystems QuantStudio 6 Flex Real-Time PCR machine using the same program as the manual method. The index PCR was kept at 10 μL to ensure a sufficient concentration after bead cleanup. The samples were cleaned using the KingFisher Apex as before. The elution plate had 25 μL of molecular-grade water, and the sample plate had 10 μL of magnetic beads. Libraries were quantified with Invitrogen Qubit 1X HS (Invitrogen, Waltham, MA, USA) and the Biotek Cytation 5 (Agilent Technologies, Santa Clara, CA, USA) instead of Quant-iT PicoGreen for the miniaturization of library quantification using the Echo 525 liquid handler, which transferred 0.6 μL of each library, while 61.4 μL of the Qubit dsDNA HS working solution buffer was added manually using a multichannel pipette. Libraries were diluted to 4 nM and pooled per preparation method.

#### Library Quantification Comparison

Libraries prepared with the manual method were quantified manually using Quant-iT PicoGreen assays, while libraries prepared with the automated method were quantified with Invitrogen Qubit 1X HS using the Echo 525 liquid handler. Because of the differences in quantification methods, the library DNA concentrations were correlated through linear regression.

#### Sequencing

All samples were first sequenced on a single 250 PE v2 MiSeq run (Illumina, San Diego, CA, USA) with a 10% PhiX spike-in. In addition, samples prepared with the Echo 525 acoustic liquid handler were also sequenced on a NextSeq 1000 (Illumina, San Diego, CA, USA) using a P1 standard cartridge with a 40% PhiX spike-in.

#### Data analysis

All bioinformatic processing and statistical analyses were performed in R (v. 4.1.3). ^27^ Sequences were assessed for quality, and ASVs were generated using DADA2 (v. 1.26.0).^28^ Low-quality reads, chimeras, chloroplast, mitochondrial, and Eukaryota reads were removed. Read counts for each method were tracked throughout sequence processing, and total read depth, average read depth, as well as high and low read depths were calculated. Contaminating reads were removed using the SCRuB package v. 0.0.1, which uses the negative and positive control samples to statistically estimate contaminants and remove them.^29^ The Phyloseq package v. 1.38.0^30^ was used for further analysis. All figures were made using the ggplot2 package^31^ in R.

Bacterial community structure was primarily analyzed using variance-stabilizing transformations on the ASV read count matrix data (DESeq2^32^ v. 1.34.0). Variance-stabilizing transformations normalize the count data and stabilize the variance, after which Euclidean distance calculations can be applied to measure community dissimilarity between samples.^33^ Secondly, to test if the large read depth disparities created by the different sequencing platforms affected community structure, we also rarefied the ASV read count matrix to 10,000 reads using the *rarefy_even_depth* function in the Phyloseq^30^ package. Following standard practice after rarefaction,^33^ a Hellinger transformation was applied to the rarefied ASV matrix to normalize the data and reduce the influence of highly abundant taxa. Bray-Curtis distances were calculated using the Vegan package (v. 2.6.4)^35^ to measure community dissimilarity. Community structure was visualized through principal coordinate analysis (PCoA) of the non-rarefied and rarefied count data. To determine significant factors affecting bacterial community structure, PERMANOVA was performed on the distance matrixes with treatment and total sequencing reads as interaction terms in the model (Vegan^35^).

Beta diversity dispersions of both variance-stabilized and Hellinger-transformed distance matrices were measured (Vegan^35^) to understand the homogeneity of dispersions based on sequencing platform and library preparation method. A Tukey HSD test was performed to determine differences in beta diversity dispersions between treatments. Additionally, to better identify differences between treatments resulting from the PERMANOVA, a comparison of beta diversity dissimilarity means both within and between treatments was performed using the *phyloseq_group_dissimilarity* function from the metagMisc package (v. 0.5.0).^36^ An ANOVA was performed on the dissimilarity values to determine the effect of sequencing platform and library preparation method, followed by a Tukey HSD post hoc test to identify differences in treatment dissimilarity means. In all instances of Tukey’s HSD test, p-values were adjusted using the Bonferroni correction.

Alpha diversity was compared between automated and manual library preparation methods sequenced on the Illumina MiSeq. Shannon diversity was calculated from the ASV read count matrix (Vegan^35^), and then correlated using a linear regression model to determine their similarities.

### Results

#### Design of a miniaturized, automated method for library preparation

We developed an automated protocol for two-step amplicon library preparation utilizing an acoustic liquid handler and an automated magnetic bead cleanup protocol. The acoustic liquid handler was incorporated into both PCR steps as well as library quantification. In both PCR stages, the PCR reaction volume was reduced five-fold (from 25 μL to 5 μL and from 50 μL to 10 μL, respectively). We also developed a protocol for automated magnetic bead cleanup using the KingFisher Apex to automate the small reaction volume bead cleanup after each PCR step (Supp. Fig. 2). Lastly, the library quantification step was also miniaturized from 200 μL (PicoGreen) to 62 μL (Qubit) with the acoustic liquid handler.

#### Comparison of sample data

The total raw read count of the automated method (4,499,351) was higher than that of the manual method (3,905,945) for samples sequenced on the Illumina MiSeq (Supp. Fig. 3). Average sample depth was 46,868 and 40,687 reads for the automated and manual methods, respectively. The sample depth ranged from 107 to 128,360 for the automated method and 2,700 to 104,444 for the manual method before filtering. After all filtering steps, there was a total of 3,456,218 reads for the automated method, and a total of 3,059,429 reads for the manual method, indicating 76.82% and 78.33% of reads were retained, respectively. The raw read count for the NextSeq was 12,656,175, with an average read depth of 131,835 and a range of 1,089 to 345,522 reads. After all filtering steps, the total read count was 9,867,362, indicating 77.96% of reads were retained.

Samples sequenced using either library preparation method with the Illumina MiSeq had more similar read depth distributions when compared to those sequenced with the Illumina NextSeq – suggesting sequencing technology had a greater effect on read depth skewness between samples (Fig. 2A). However, this could also be due to individual library preparation, sample pooling, and library quantification. When comparing the samples sequenced by the Illumina MiSeq, those prepared with the automated method generally had a higher read depth than those prepared with the manual method (Fig. 2B). However, this could be due to the final concentration of pooled libraries and human errors associated with manual pooling and loading of the sequencing cartridge. Samples prepared manually had a lower read depth variance (5.1 x 10^8^) but a higher level of skewness (0.63). Whereas samples prepared with the automated method had a higher read depth variance (8.1 x 10^8^) but a lower skewness (0.43; Fig. 2B).

**Figure 2.**
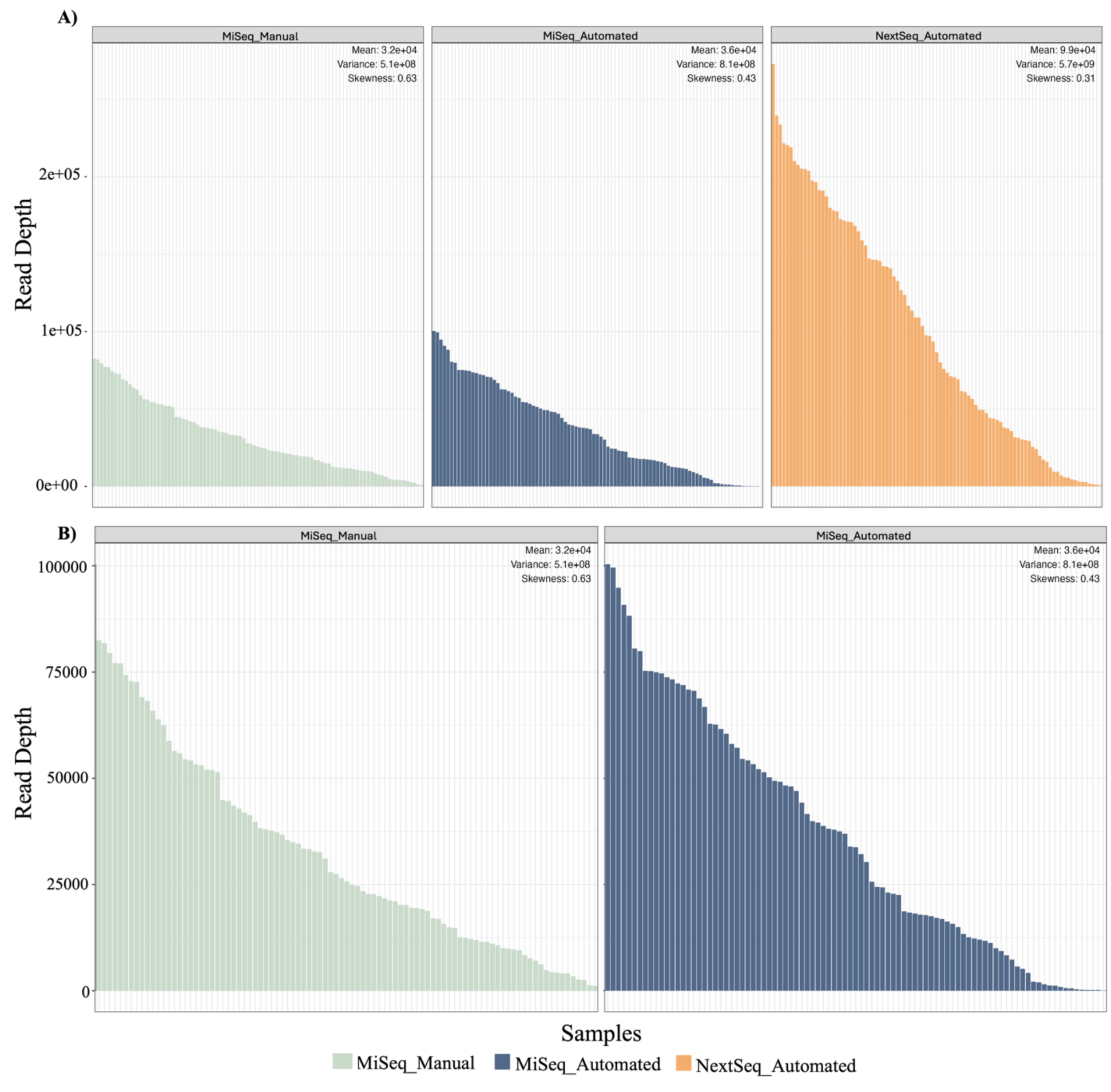
Automated library preparation has generally led to an increase in read depth. **A)** The distribution of each sample’s read depth across sequencing platform and library preparation method is shown in the above histogram plots. **B)** The distribution of read depth of samples sequenced on the Illumina MiSeq only. The mean, variance, and skewness of each was calculated. Values for these are given in the top right of each figure.

The Shannon diversity of samples was highly correlated (R^2^_adj_ 0.74; Fig. 3) indicating the comparability of the manual and automated library preparation methods in taxa detection.

**Figure 3.**
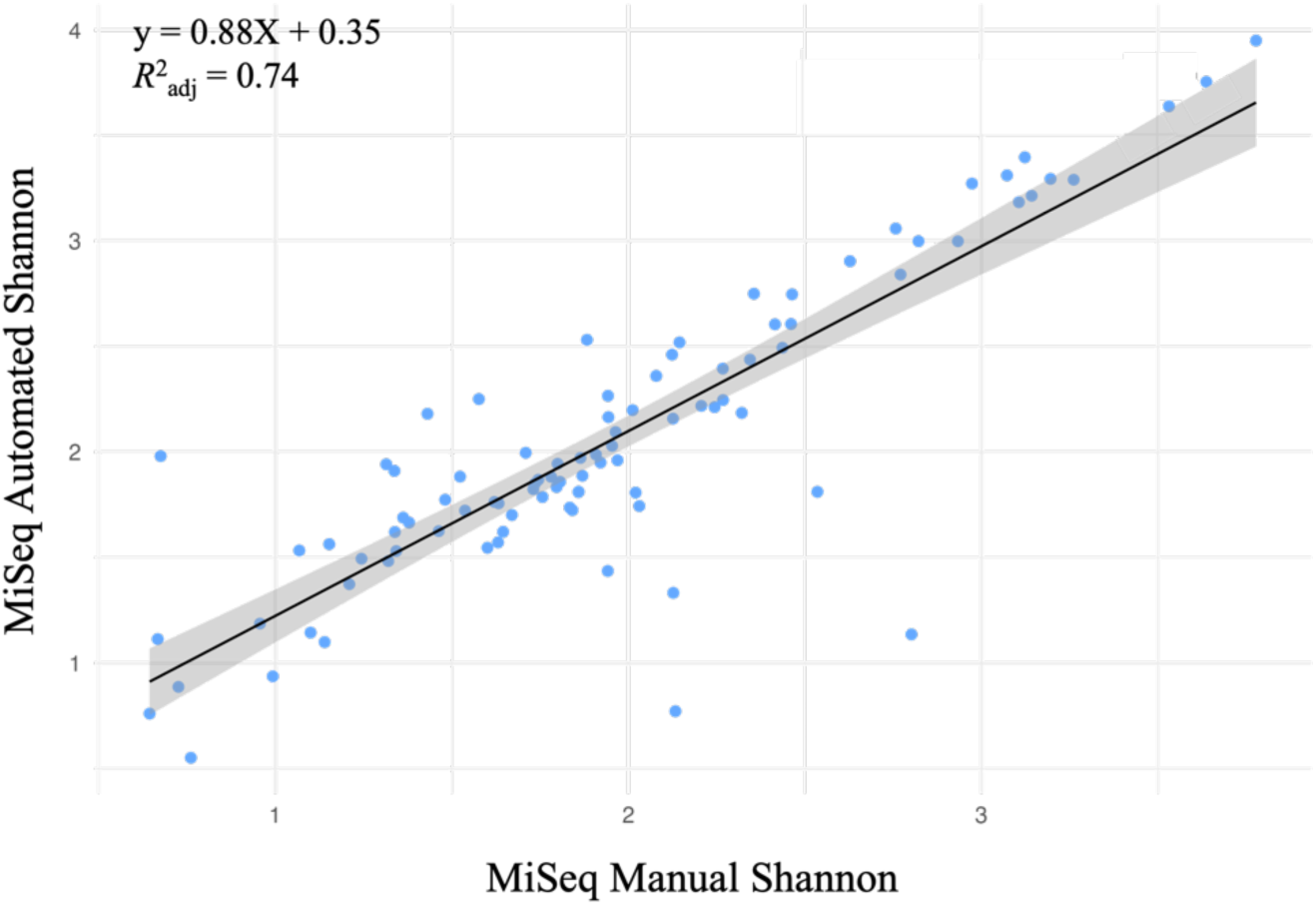
Library preparation methods have similar bacterial alpha diversity metrics. Correlation of Shannon alpha diversity values of automated and manual library preparation methods sequenced on the Illumina MiSeq. Data points represent individual samples.

Bacterial community structure varied by sequencing platform depending on library preparation method (interaction p-value 0.001***; Fig. 4A). Additionally, bacterial community structure also differed by sample read depth depending on sequencing platform and library preparation method (interaction p-value 0.001***). Samples sequenced on the Illumina NextSeq showed a greater beta-dispersion in community composition compared to both MiSeq preparation methods (all p-value <0.001***; Supp. Fig. 5A), which was likely a result of the increased read depth compared to the MiSeq (Fig. 4B). Additionally, samples sequenced on the NextSeq had higher average beta diversity dissimilarities alone and compared to both MiSeq methods. The average dissimilarities for the automated MiSeq method were also higher than the manual MiSeq method. However, the comparison between both MiSeq methods was not different from the within-treatment comparison of both methods (Supp. Fig. 6A).

**Figure 4.**
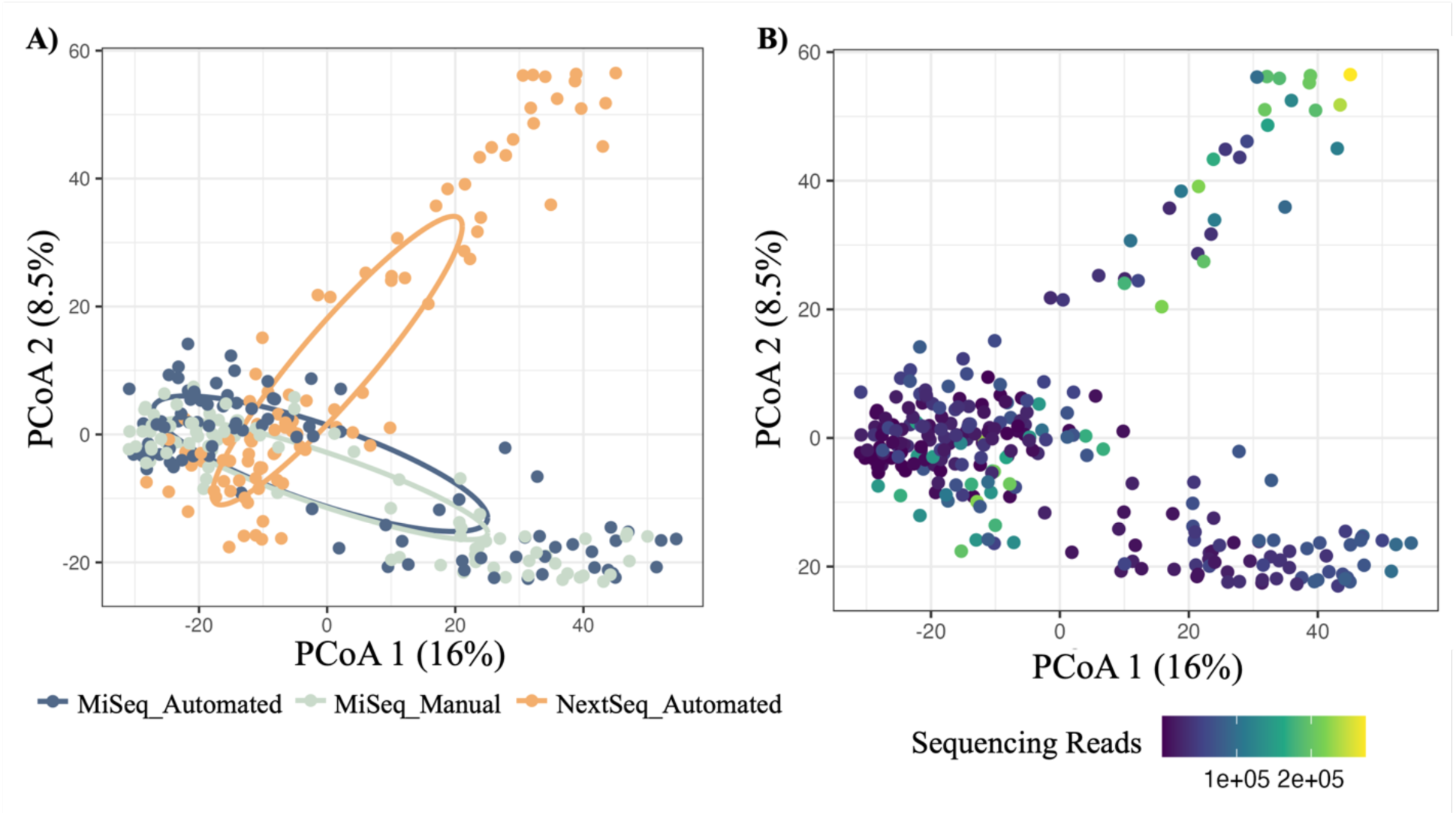
Bacterial community structure varies by sequencing platform but not library preparation method, due to sequencing depth differences. PCoA plots showing community structure differences of variance-stabilized data with Euclidean distances by **A)** sequencing platform and library preparation method, and **B)** sequencing depth. A lighter color represents a greater number of reads. Each point represents a single sample. P-values: sequencing platform and library preparation method x read depth interaction (0.001***), sequencing platform and library preparation method (0.001***), read depth (0.001***).

Rarefaction curves were generated (Supp. Fig. 4) and determined that 10,000 reads were the required per sample to capture the majority of taxa present while limiting the number of samples removed due to low read depth. Although samples visually appeared to have small differences in communities between samples sequenced on the MiSeq and NextSeq after rarefaction, there were still significant differences in communities depending on the sequencing platform and library preparation method (p-value: 0.001***; Fig. 5). However, beta diversity dispersion was not significantly different between any of the sequencing platforms and library preparation methods (TukeyHSD p-adjusted values range from 0.873 to 0.993; Supp. Fig. 5B). Comparison of the average beta diversity dissimilarities revealed that while rarefaction has removed the effect of library preparation method for samples sequenced on the MiSeq, it did not completely remove the NextSeq platform effect on community composition (Supp. Fig. 6B).

**Figure 5.**
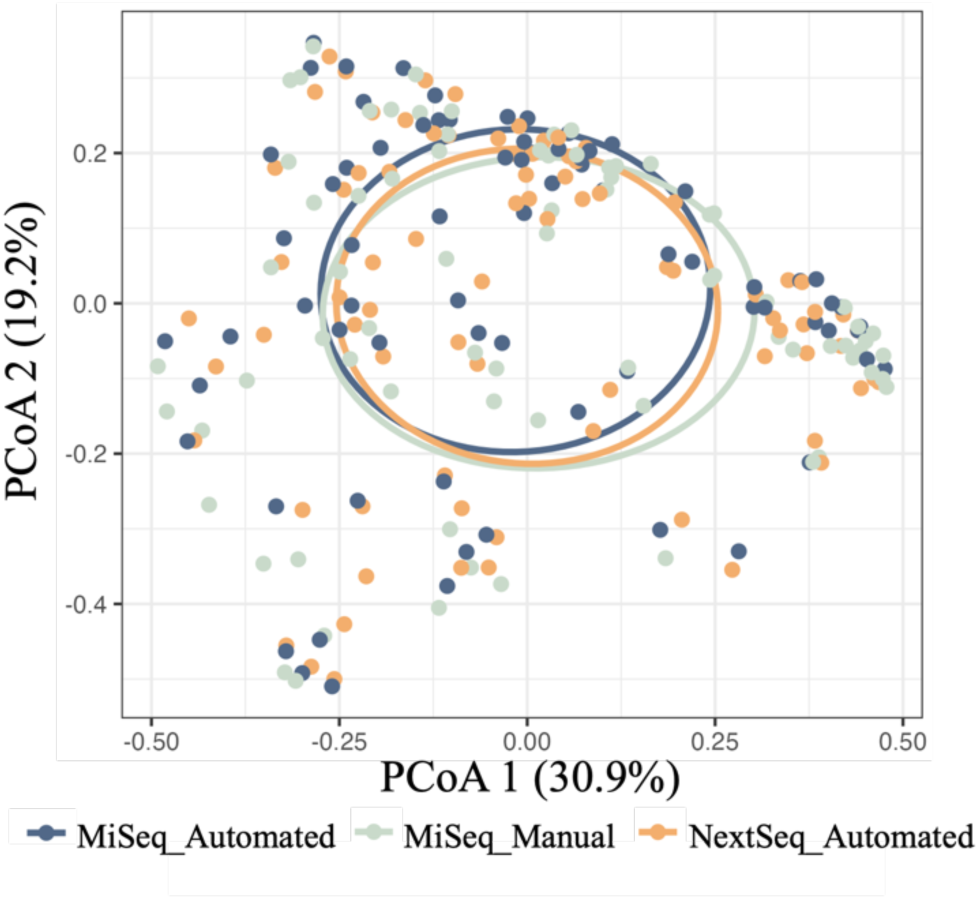
Rarefaction of sequencing results nearly eliminates the differences in community structure due to greater sequencing depth. PCoA plot of sequencing results after rarefying each sample to 10,000 reads. P-value: sequencing platform and library preparation method (0.001***).

#### Comparison of quantification

After preparation, the PicoGreen and Qubit DNA quantification methods were compared between manual and automated library preparation methods, respectively. Similar to a previous test comparing both quantification methods,^37^ the automated Qubit quantification method was highly correlated with the manual PicoGreen method (Adj. R^2^ = 0.98; Fig. 6). However, evaluation of the slope from the regression (0.83) indicated that the automated Qubit method tended to have lower sample concentrations, relative to the manual PicoGreen method – though the between-sample comparisons within a method did not vary along the range of concentrations tested.

**Figure 6.**
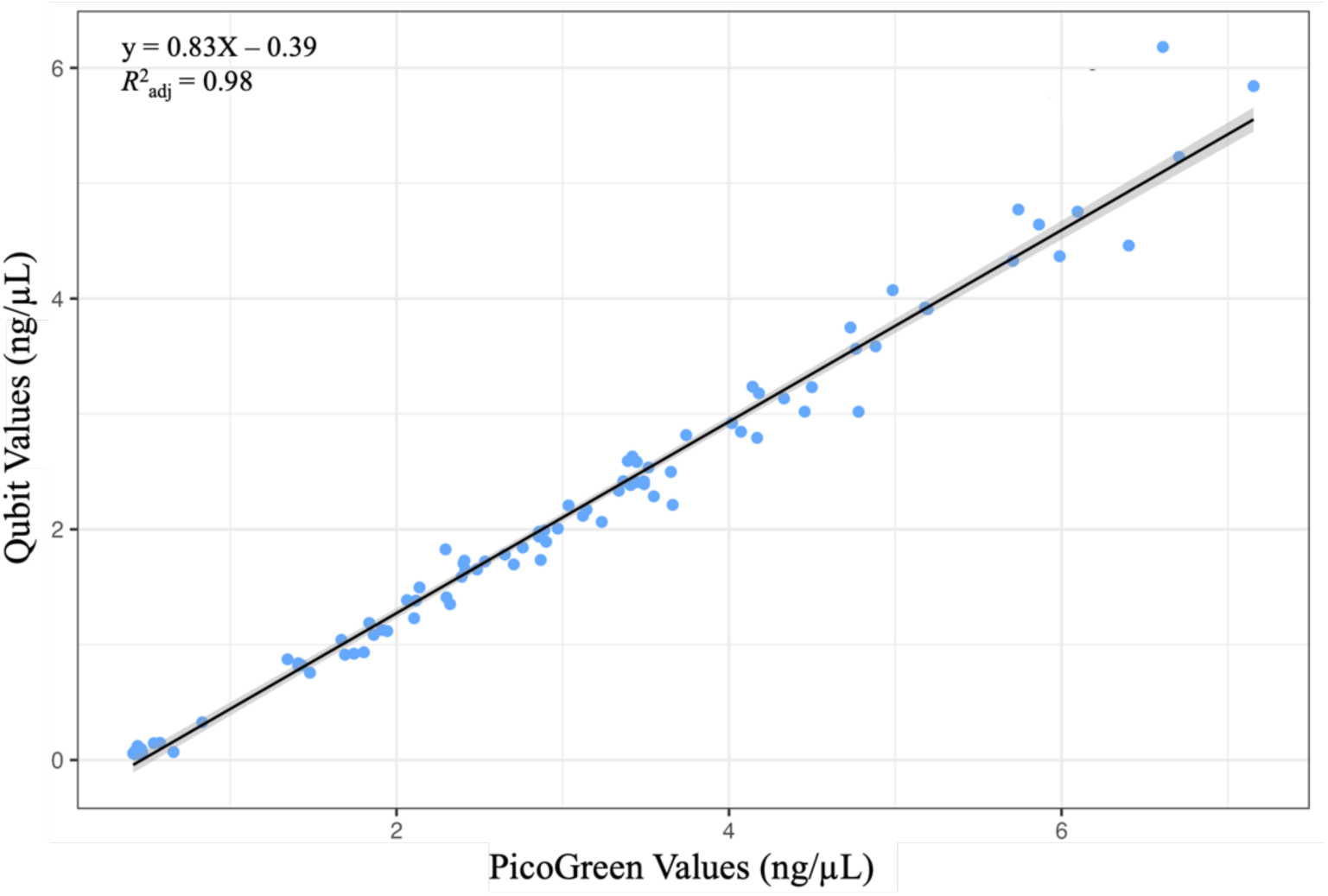
Manual PicoGreen and automated Qubit quantification methods are strongly correlated. Correlation of DNA quantification with PicoGreen (manual) and Qubit (automated). Data points represent individual samples. Quantification measured in ng/μL. Line represents linear regression model.

#### Comparison of consumable usage

The consumables were tracked for manual and automated library preparation, and the person-hours for each method were also estimated and are presented in Table 1. The development of the automated library preparation protocol led to a reduction in costs as well as person-hours (Table 1). The price reductions for each step in the library preparation were directly compared. There was a ∼50% reduction in price for both the amplicon and index PCRs using the automated method (Fig. 7). There was a ∼15-30% reduction in costs for all other steps, except for gel electrophoresis and TapeStation sample check steps – where the automated and manual method costs were comparable (Fig. 7). The cost to prepare libraries for one 96-well plate of samples using the manual method was ∼$2,200, while the automated method was reduced to ∼$1,500 (Table 1), which led to a ∼30% reduction in total costs. The total costs of consumables were tracked and totaled for each step and each method (Supp. Table 1). There was a ∼50% reduction in person-hours to prepare libraries with the automated method (6.5 hours) compared to the manual method (12.42 hours). Thus, the automated library preparation protocol decreased both the costs and time to prepare libraries.

**Figure 7.**
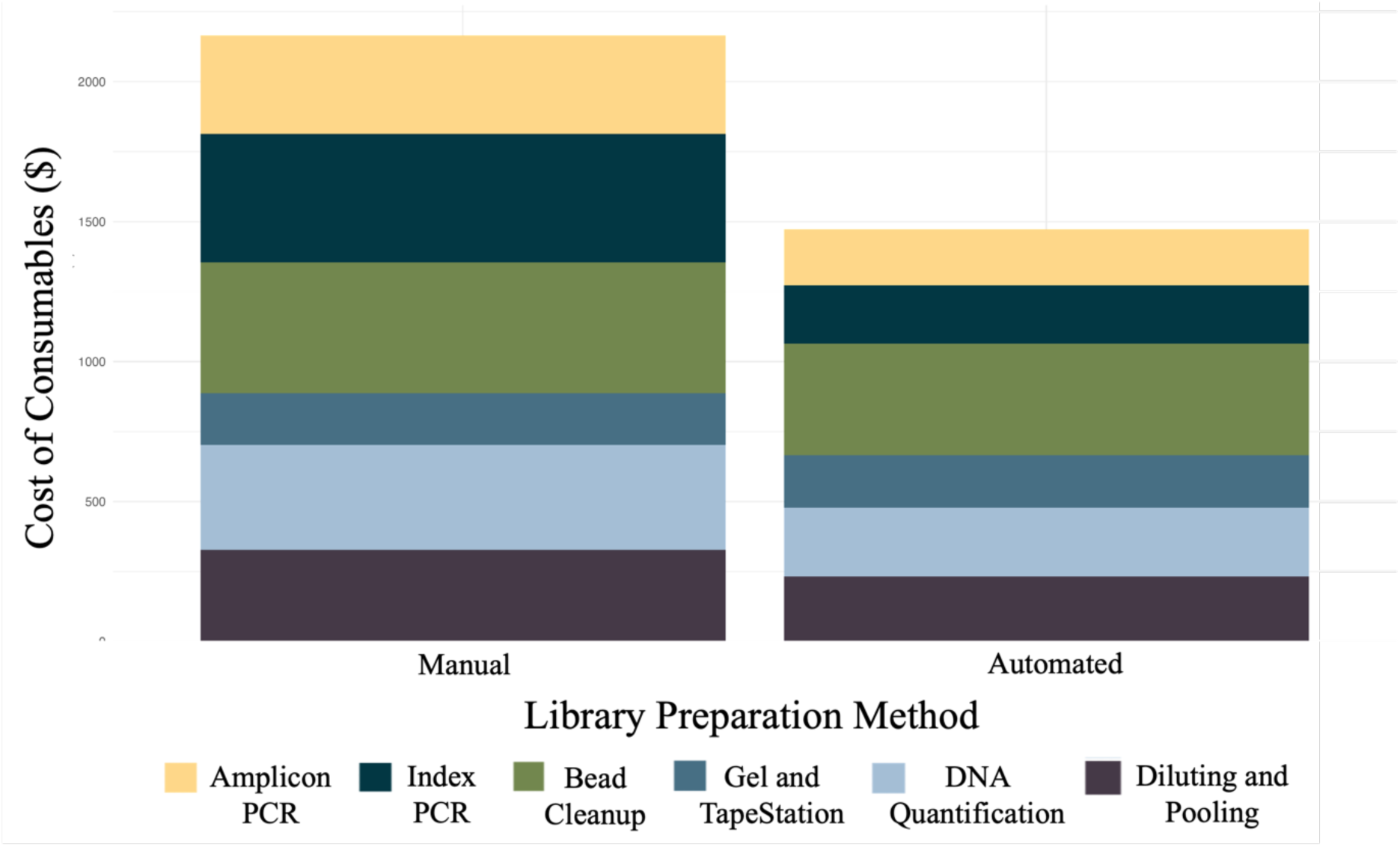
The automated method has decreased the cost of library preparation. The cost of consumables and reagents for each method, colored by the totals for the library preparation step.

**Table 1.**
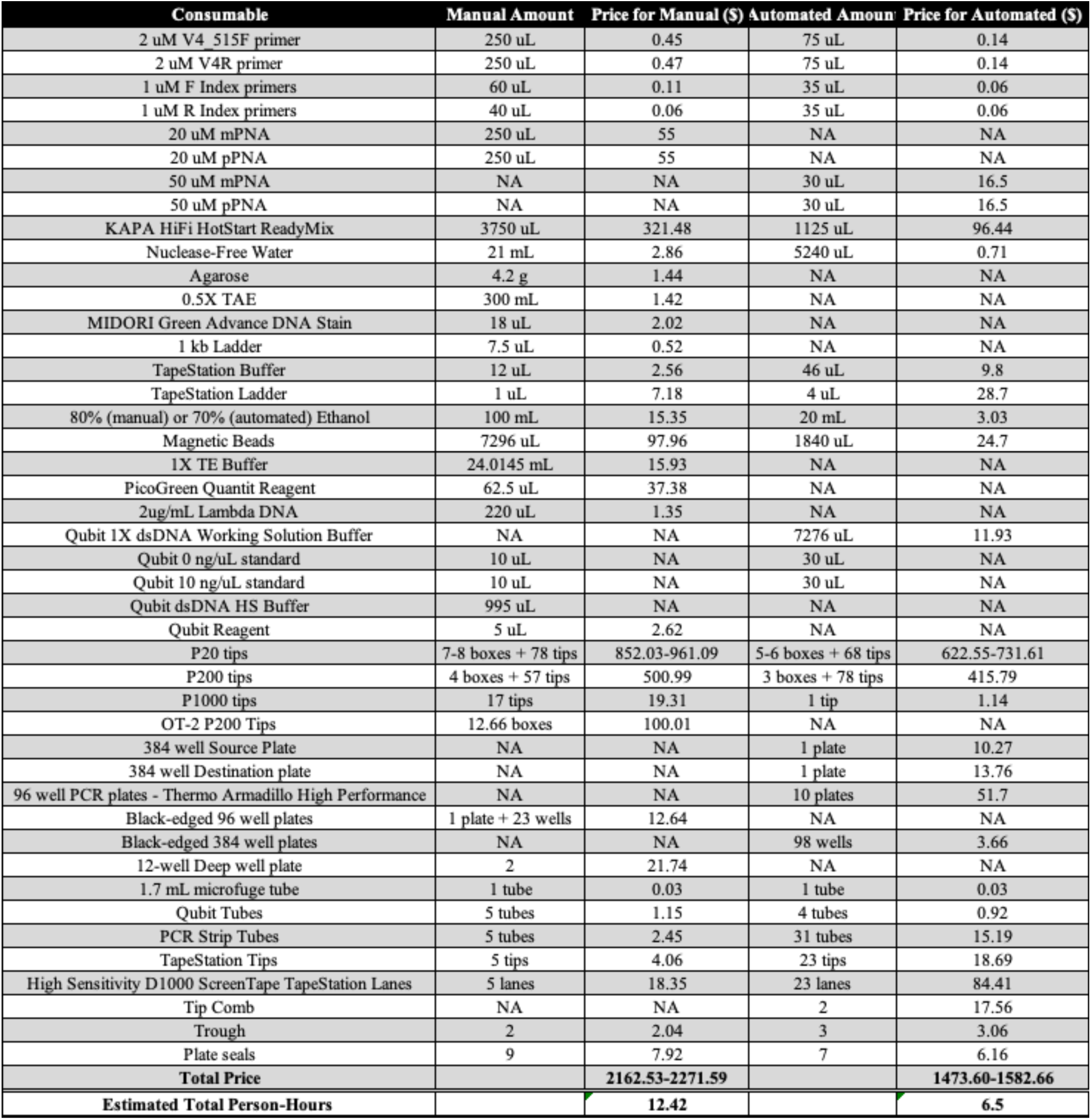
The automated library preparation method has decreased the time and cost of library preparation compared to the manual method. Table comparing consumables and reagents used for both manual and automated library preparation. Volumes or number of items used are totaled for each reagent or consumable. Prices for each consumable are given per preparation method. Estimated person-hour totals are given per preparation

| Consumable | Manual Amount | Price for Manual (\$) | Automated Amount | Price for Automated (\$) |
| --- | --- | --- | --- | --- |
| 2 uM V4 515F primer | 250 uL | 0.45 | 75 uL | 0.14 |
| 2 uM V4R primer | 250 uL | 0.47 | 75 uL | 0.14 |
| 1 uM F Index primers | 60 uL | 0.11 | 35 uL | 0.06 |
| 1 uM R Index primers | 40 uL | 0.06 | 35 uL | 0.06 |
| 20 uM mPNA | 250 uL | 55 | NA | NA |
| 20 uM pPNA | 250 uL | 55 | NA | NA |
| 50 uM mPNA | NA | NA | 30 uL | 16.5 |
| 50 uM pPNA | NA | NA | 30 uL | 16.5 |
| KAPA HiFi HotStart ReadyMix | 3750 uL | 321.48 | 1125 uL | 96.44 |
| Nuclease-Free Water | 21 mL | 2.86 | 5240 uL | 0.71 |
| Agarose | 4.2 g | 1.44 | NA | NA |
| 0.5X TAE | 300 mL | 1.42 | NA | NA |
| MIDORI Green Advance DNA Stain | 18 uL | 2.02 | NA | NA |
| 1 kb Ladder | 7.5 uL | 0.52 | NA | NA |
| TapeStation Buffer | 12 uL | 2.56 | 46 uL | 9.8 |
| TapeStation Ladder | 1 uL | 7.18 | 4 uL | 28.7 |
| 80% (manual) or 70% (automated) Ethanol | 100 mL | 15.35 | 20 mL | 3.03 |
| Magnetic Beads | 7296 uL | 97.96 | 1840 uL | 24.7 |
| 1X TE Buffer | 24.0145 mL | 15.93 | NA | NA |
| PicoGreen Quantit Reagent | 62.5 uL | 37.38 | NA | NA |
| 2ug/mL Lambda DNA | 220 uL | 1.35 | NA | NA |
| Qubit 1X dsDNA Working Solution Buffer | NA | NA | 7276 uL | 11.93 |
| Qubit 0 ng/uL standard | 10 uL | NA | 30 uL | NA |
| Qubit 10 ng/uL standard | 10 uL | NA | 30 uL | NA |
| Qubit dsDNA HS Buffer | 995 uL | NA | NA | NA |
| Qubit Reagent | 5 uL | 2.62 | NA | NA |
| P20 tips | 7-8 boxes + 78 tips | 852.03-961.09 | 5-6 boxes + 68 tips | 622.55-731.61 |
| P200 tips | 4 boxes + 57 tips | 500.99 | 3 boxes + 78 tips | 415.79 |
| P1000 tips | 17 tips | 19.31 | 1 tip | 1.14 |
| OT-2 P200 Tips | 12.66 boxes | 100.01 | NA | NA |
| 384 well Source Plate | NA | NA | 1 plate | 10.27 |
| 384 well Destination plate | NA | NA | 1 plate | 13.76 |
| 96 well PCR plates - Thermo Armadillo High Performance | NA | NA | 10 plates | 51.7 |
| Black-edged 96 well plates | 1 plate + 23 wells | 12.64 | NA | NA |
| Black-edged 384 well plates | NA | NA | 98 wells | 3.66 |
| 12-well Deep well plate | 2 | 21.74 | NA | NA |
| 1.7 mL microfuge tube | 1 tube | 0.03 | 1 tube | 0.03 |
| Qubit Tubes | 5 tubes | 1.15 | 4 tubes | 0.92 |
| PCR Strip Tubes | 5 tubes | 2.45 | 31 tubes | 15.19 |
| TapeStation Tips | 5 tips | 4.06 | 23 tips | 18.69 |
| High Sensitivity D1000 ScreenTape TapeStation Lanes | 5 lanes | 18.35 | 23 lanes | 84.41 |
| Tip Comb | NA | NA | 2 | 17.56 |
| Trough | 2 | 2.04 | 3 | 3.06 |
| Plate seals | 9 | 7.92 | 7 | 6.16 |
| **Total Price** |  | **2162.53-2271.59** |  | **1473.60-1582.66** |
| **Estimated Total Person-Hours** |  | **12.42** |  | **6.5** |

#### Discussion

The availability and reduced costs of sequencing have led to an increase in studies utilizing high-throughput sequencing technologies. However, there is still a significant expense for the preparation of samples for sequencing due to reagent and consumable costs. The incorporation of an acoustic liquid handler enabled the development of a library preparation protocol that decreased the preparation time and reaction volumes, ultimately increasing the efficiency of material use and lowering user expenses (Fig. 1, Table 1).

We compared this automated method to the standard, manual preparation. The automated library preparation method led to a higher average read depth and a more symmetrical distribution in sequencing (Fig. 2). The taxa detection and Shannon diversity of both automated and manual methods were comparable (Fig. 3). When analyzing the bacterial community compositions, we saw that while library preparation method significantly affected these communities, a majority of this effect was due to an increase in sequencing read depth (Fig. 4). After rarefaction, this effect was still significant but much lower, and there were fewer differences in community compositions (Fig. 5, Supp. Fig. 6). This could be due to the increase in read depth seen with the automated library preparation method, which would allow for a greater understanding of microbial communities including rare or low-abundance taxa.^38,39^ The automation and miniaturization reduced the time and cost of preparing samples for amplicon sequencing by ∼50% (Fig. 6, Table 1). Our cost of preparation for one 96-well plate was over $2,000 for the manual method or ∼$23 per sample. The automated method has reduced that cost to around $1,500 or ∼$16 per sample. This approach not only lowers expenses but also supports the environmental sustainability of research by limiting plastic waste.

There are several studies using acoustic liquid handlers to miniaturize and automate different types of molecular and cellular assays,^15–20^ as well as one for amplicon sequencing.^21^ In this previous report, EMP researchers miniaturized a one-step library preparation method using an acoustic liquid handler and compared it to a pipette-based liquid handler, which led to a ∼58% reduction in library preparation costs. Our method led to a ∼30% reduction in expenses due to miniaturization. However, this is likely because we used a two-step PCR process, which requires more polymerase, magnetic beads, and transfers between PCRs and bead cleanups compared to the one-step method. The two-step PCR method has several advantages, including: cheaper primers, reduced amplification bias when compared to the long adapter index primers required in the one-step method,^23^ and the option to increase sample multiplexing by introducing a second layer of barcodes during amplicon PCR.^24^ While the previous group successfully miniaturized an amplicon library preparation protocol using an acoustic liquid handler, we combined an acoustic liquid handler and an automated magnetic bead cleanup system for a two-step PCR library preparation process.

We successfully developed a method that miniaturized and reduced the costs of amplicon library preparation for microbiome studies compared to the standard, manual method. While this method was tested on the V4 region of the 16S rRNA gene, other regions of the 16S rRNA gene and the ITS region could be sequenced using the corresponding primer sets. Our automated protocol is available for further optimization and cost-saving changes. For example, since the initial test of this protocol, we have reduced tip waste by eliminating a sample transfer between 384-well and 96-well plates after PCR amplification prior to bead cleanup. This was accomplished by simply using 96-well plates throughout all steps and further reduced sample costs from $16 to $13 per sample, which will almost fully offset the costs of a sequencing cartridge. One limitation of this method is the necessity to use specific equipment, including the acoustic liquid handler and automated magnetic bead cleanup robot.

#### Conclusions

Due to the need to reduce the bottleneck of library preparation in amplicon sequencing, we successfully developed a miniaturized and automated method utilizing an acoustic liquid handler paired with automated magnetic bead cleanup. The ability of the liquid handler to transfer small volumes of liquid increased efficiency in consumables and reagent usage and reduced time for completion compared to the manual method. A major benefit of the automated preparation method is the reduction in consumable costs and person-hours (Fig. 7, Table 1) with no differences in bacterial communities from the MiSeq (Fig. 5, Supp. Fig. 5). The increased consumable efficiency and reduced time to prepare libraries using this method will lead to significant cost savings to prepare hundreds of samples in future microbiome studies.

## 8. Author statements

### 8.1 Author contributions

Conceptualization, B.R.B., E.L.E., and B.A.G.; Methodology, B.R.B., E.L.E. and B.A.G.; Validation, B.R.B.; Writing – original draft preparation, B.R.B.; Writing – review and editing, B.R.B., B.K.W., T.B. and B.A.G.; Supervision, T.B. and B.A.G.

## 8.4 Acknowledgements

We thank the following people and institutions for help with fieldwork, sample collection, and sample processing: Patrick Gross, Abraham Hangamaisho, Abbeah R. Navasca, and Diel D. Velasco (North Dakota State University), Dr. Mark Sorrells and James Tanaka (Cornell University), and Dr. Ruth Dill-Macky and Kelsey Hyland (University of Minnesota). We thank Megan Ramsett (North Dakota State University) and the Dr. Thomas Glass Biotech Innovation Core at North Dakota State University for providing support in library preparation and amplicon sequencing specifically Scott Hoselton and Kaycie Schmidt. We also thank Martha Vaughan (USDA-ARS) for her help with the approval and review of this paper. Project funding was provided by the State Board of Agricultural Research and Education (SBARE). This work was supported in part by the U.S. Department of Agriculture (USDA), Agricultural Research Service. Mention of trade names or commercial products in this publication is solely for the purpose of providing specific information and does not imply recommendation or endorsement by the USDA. The USDA is an equal opportunity provider and employer.

## Supplementary

**Supplementary Figure 1.**
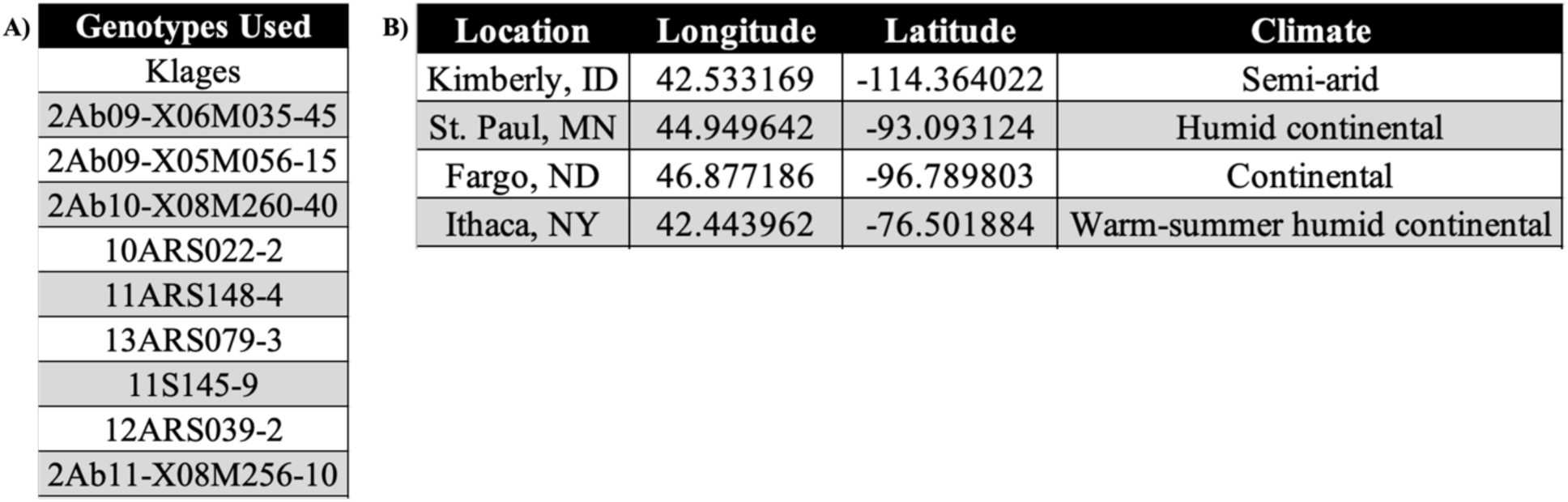
**A)** 10 specific barley genotypes from the spring malt barley training population. **B)** Longitude, latitude, and climate type of the four FHB-misted nursery locations.

**Supplementary Figure 2.**
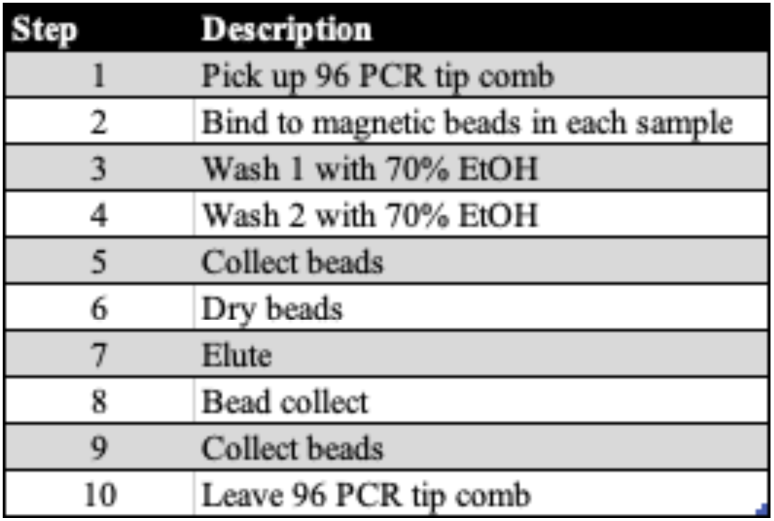
Steps for small volume magnetic bead cleanup program for the KingFisher Apex.

**Supplementary Figure 3.**
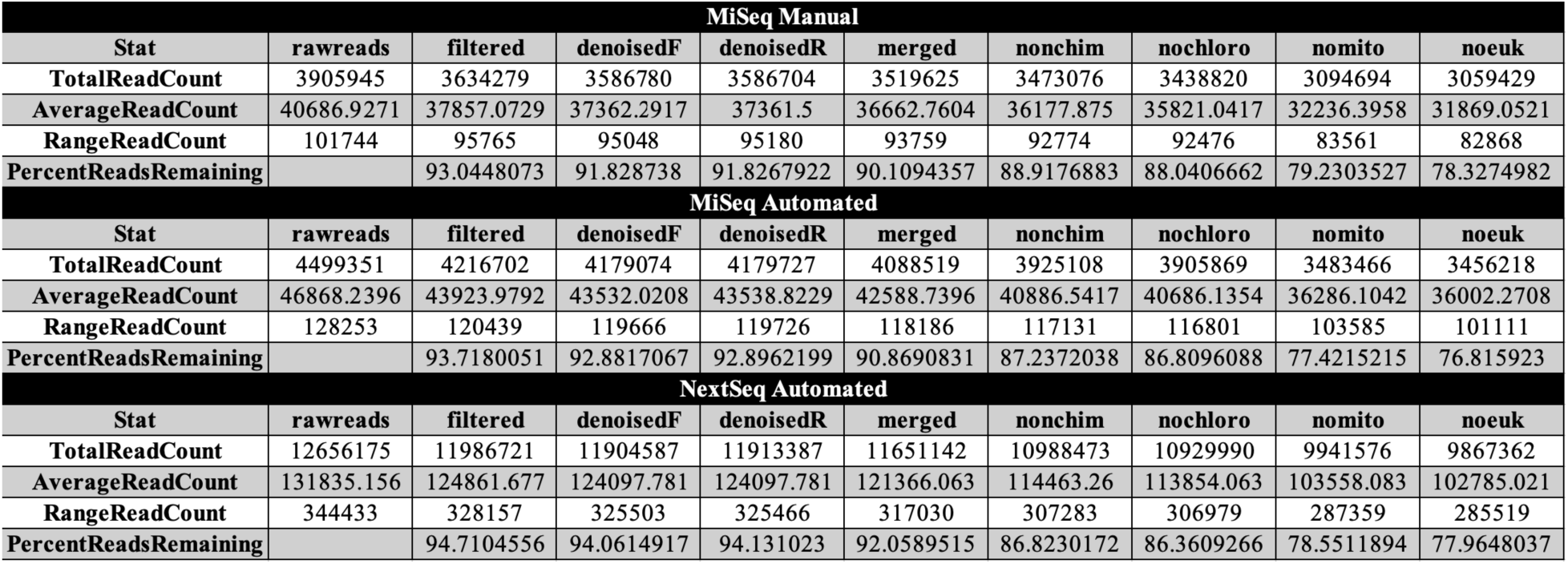
Tracking read count, the average read counts, the range of read counts, and the percent of reads remaining after each filtering step in DADA2.

**Supplementary Figure 4.**
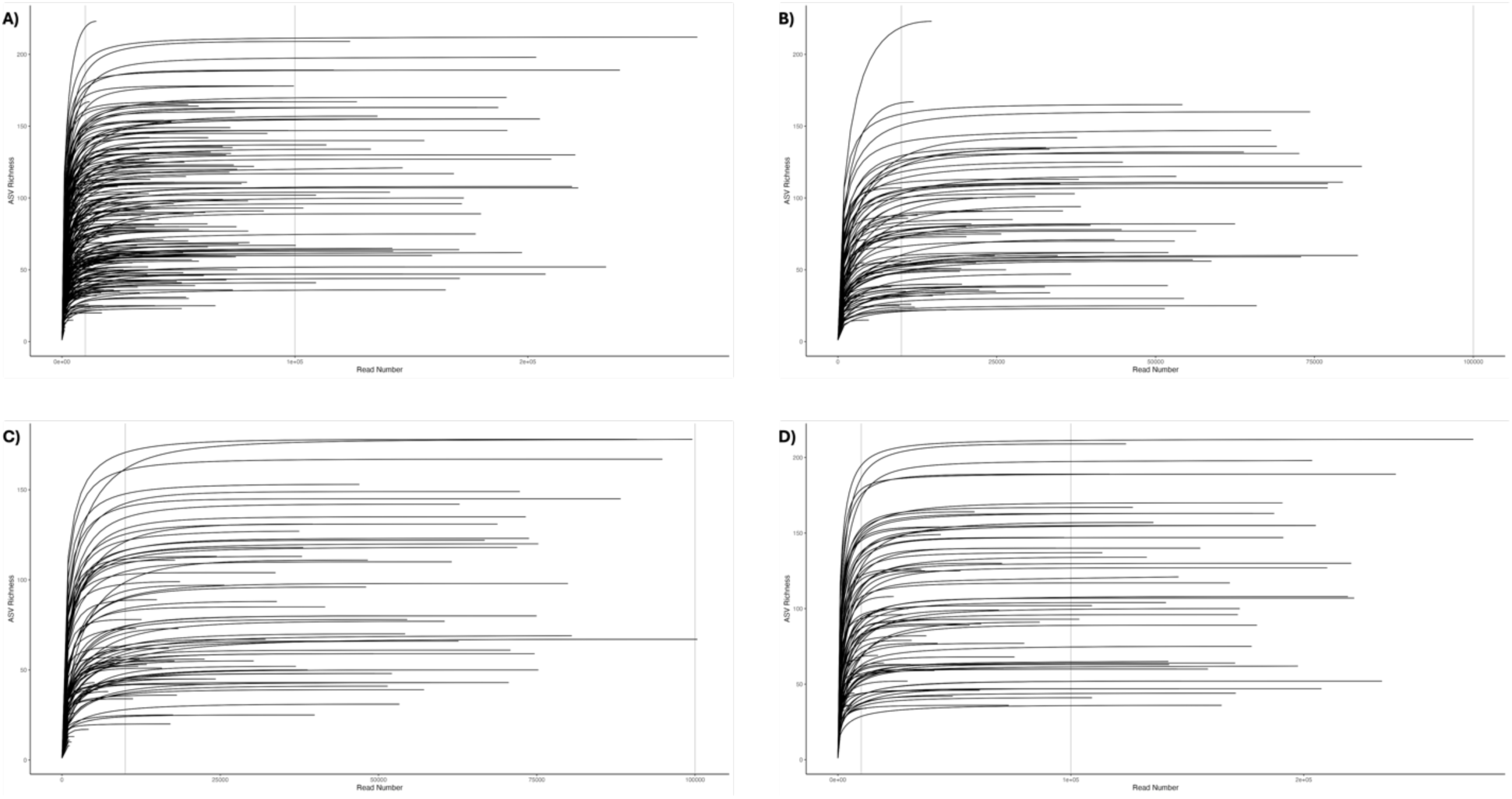
Rarefaction curves of each sample showing read depth along the x-axis and ASV richness along the y-axis. The 10,000 and 100,000 read-depths are denoted with vertical grey lines. **A)** Rarefaction curves of all samples across each sequencing platform and library preparation method. **B)** Rarefaction curves of samples prepared manually and sequenced by the MiSeq. **C)** Rarefaction curves of samples prepared with the acoustic liquid handler and sequenced by the MiSeq. **D)** Rarefaction curves of samples prepared with the acoustic liquid handler and sequenced by the NextSeq.

**Supplementary Figure 5.**
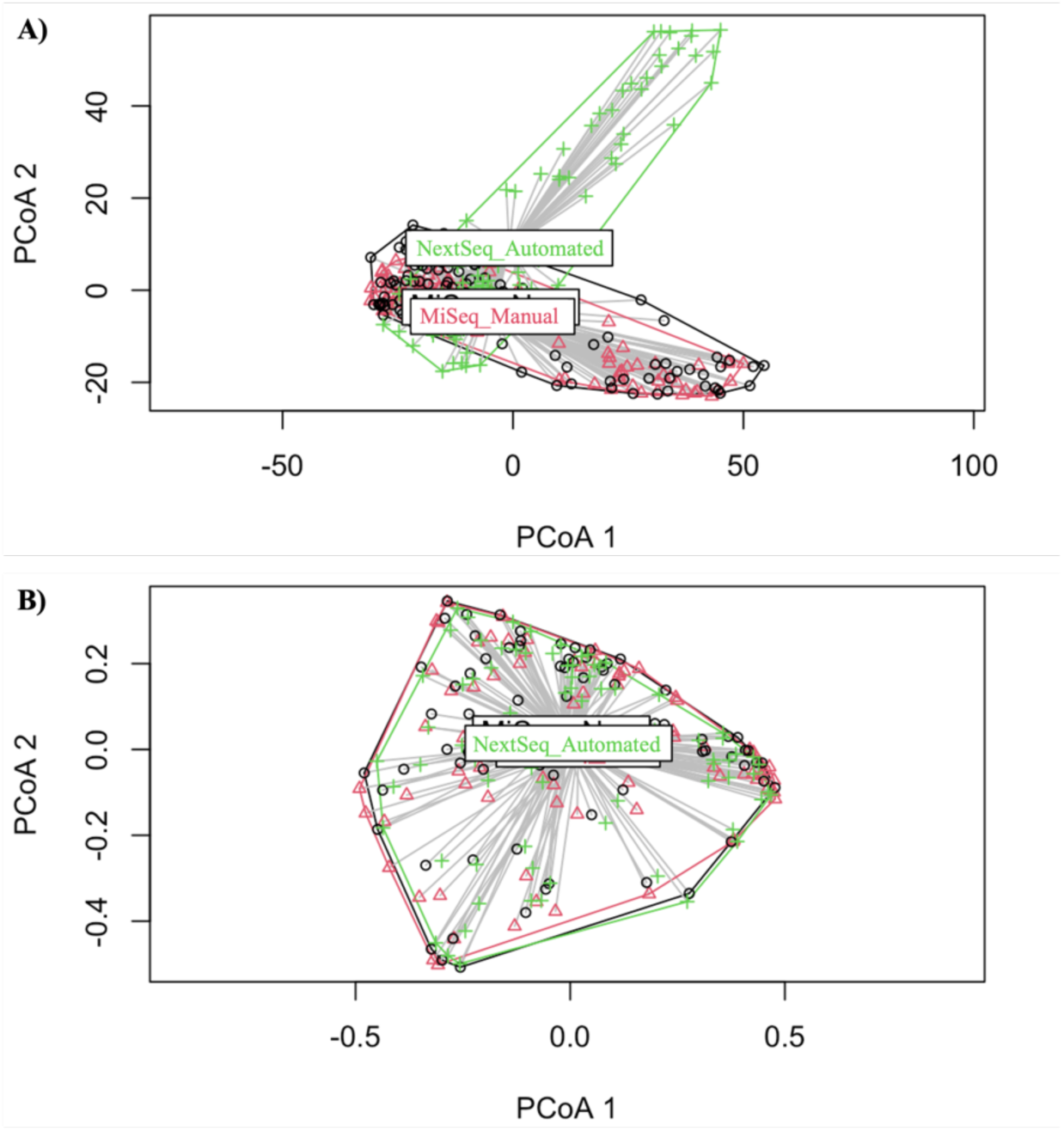
Beta diversity dispersion of sequencing platform and library preparation methods. Each point represents a single sample. Significance was tested using a Tukey HSD test. **A)** Dispersion of samples before rarefaction. P-adjusted values: MiSeq Manual x MiSeq Automated (0.217), NextSeq Automated x MiSeq Manual (<0.001***), NextSeq Automated x MiSeq Automated (<0.001***). **B)** Dispersion of samples after rarefaction. P-adjusted values: MiSeq Manual x MiSeq Automated (0.873), NextSeq Automated x MiSeq Manual (0.919), NextSeq Automated x MiSeq Automated (0.993).

**Supplementary Figure 6.**
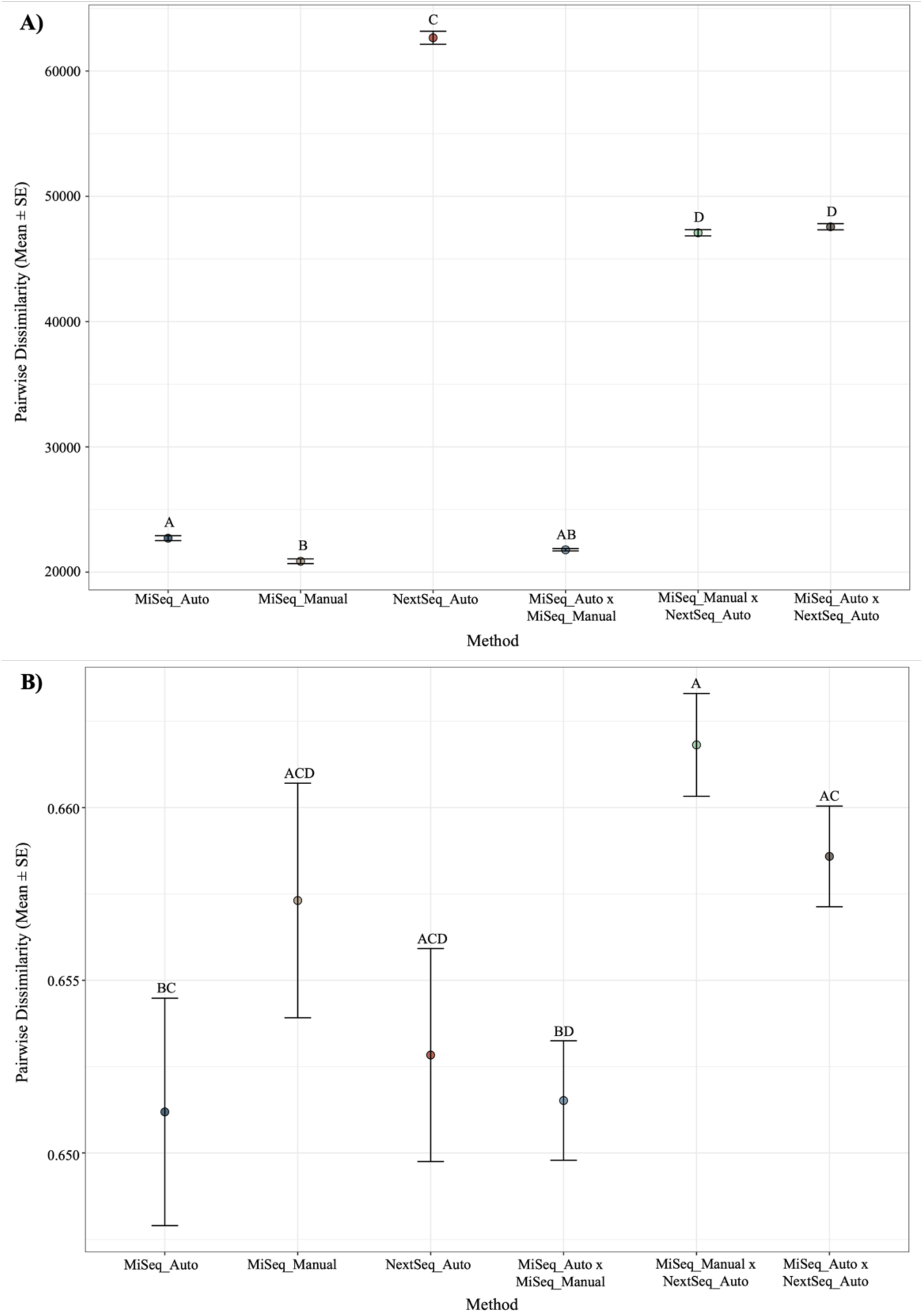
Comparison of beta diversity dissimilarity means for each sequencing platform and library preparation method and their pairwise comparisons. Significance was determined with a Tukey HSD test. **A)** Comparison of means before rarefaction using Euclidean distances. **B)** Comparison of means after rarefaction using Bray-Curtis distances.

**Supplementary Table 1.**
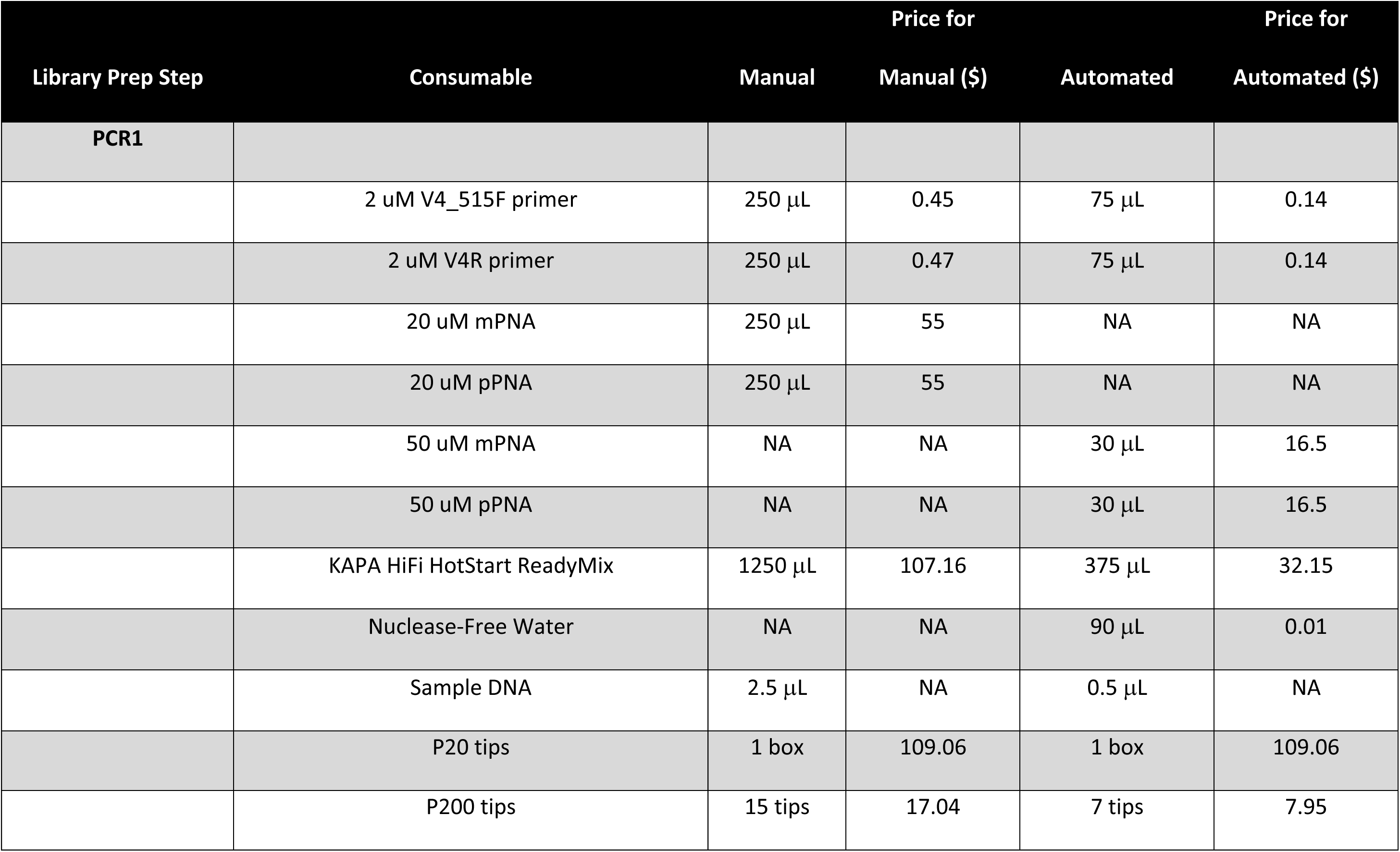
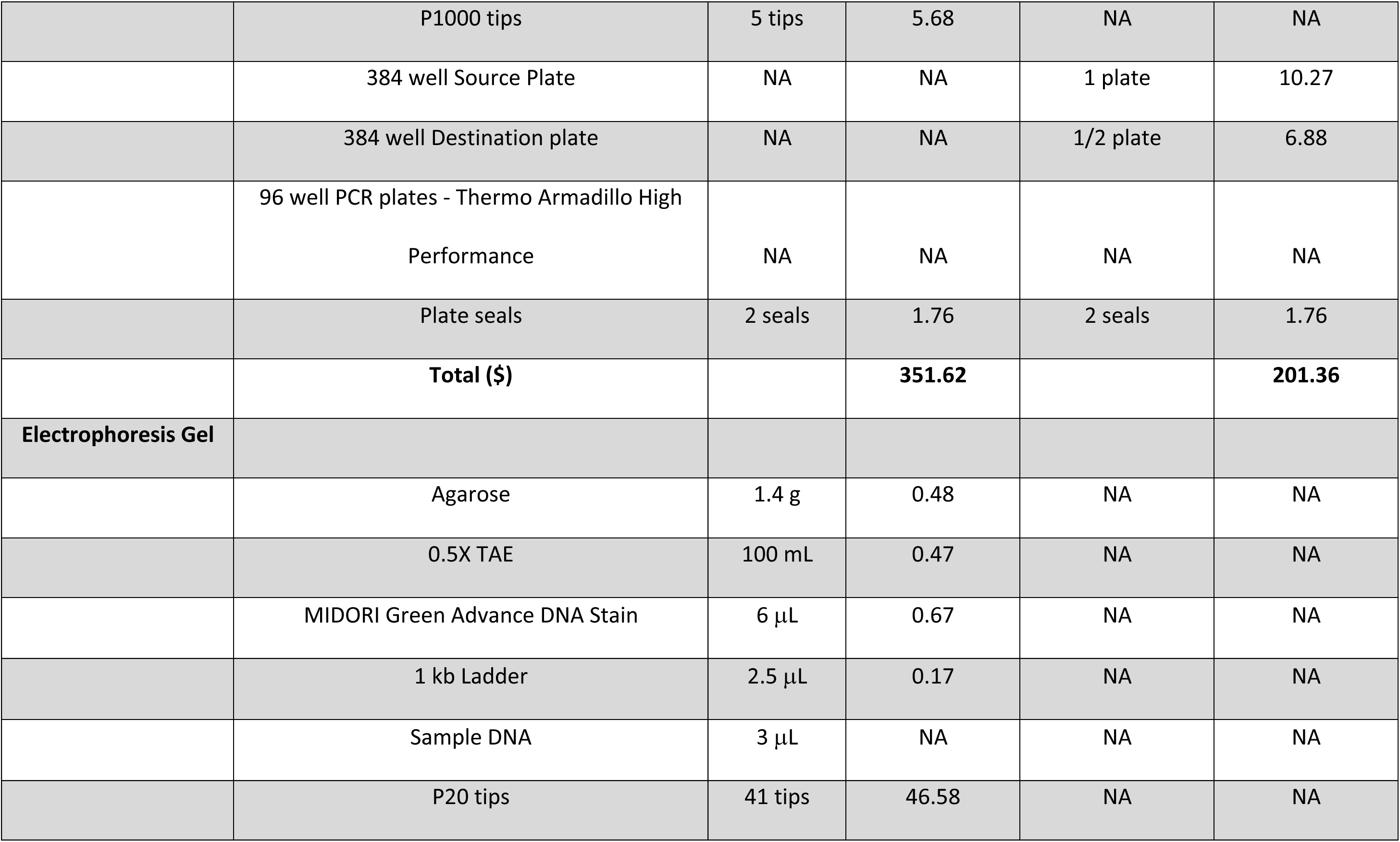
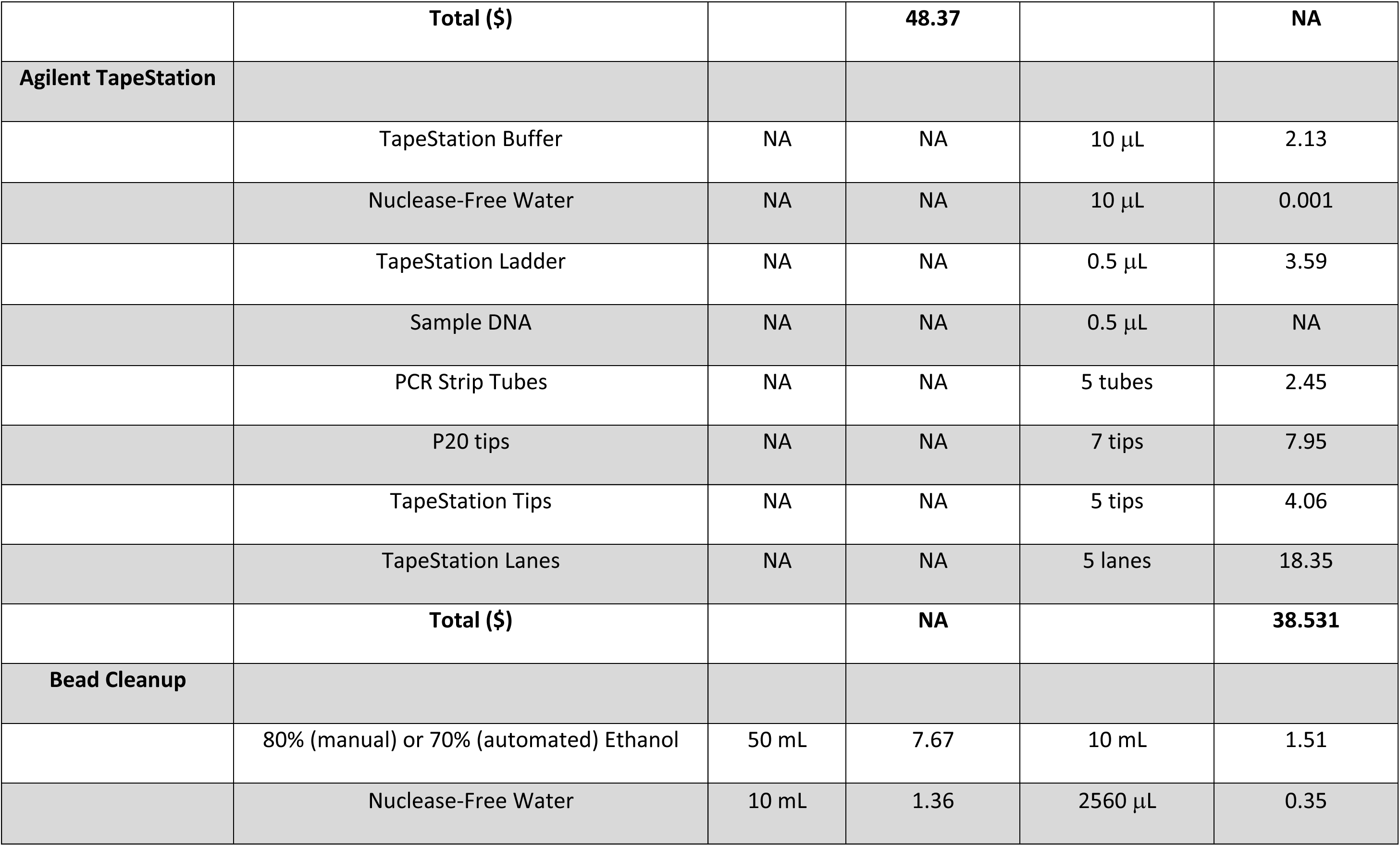
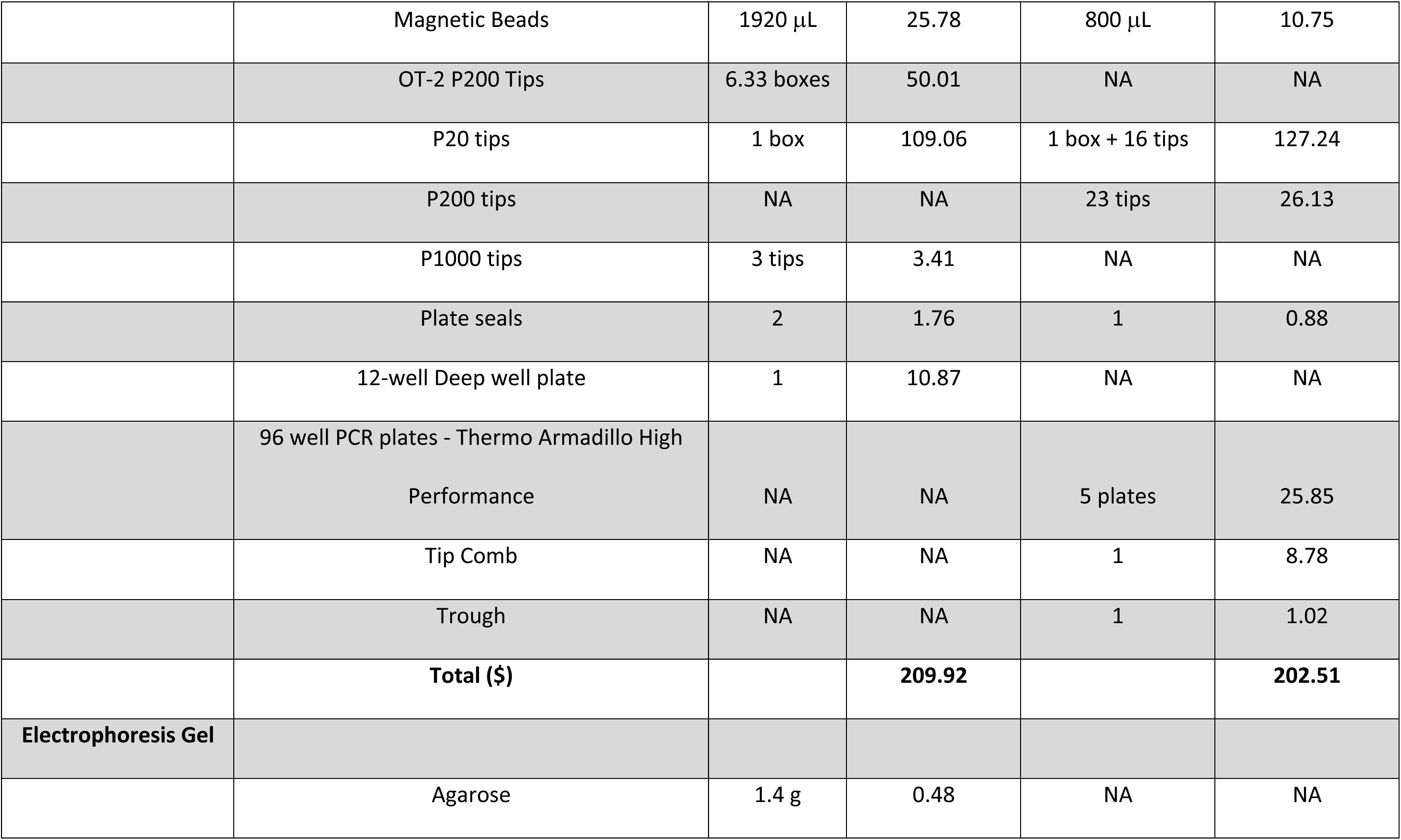
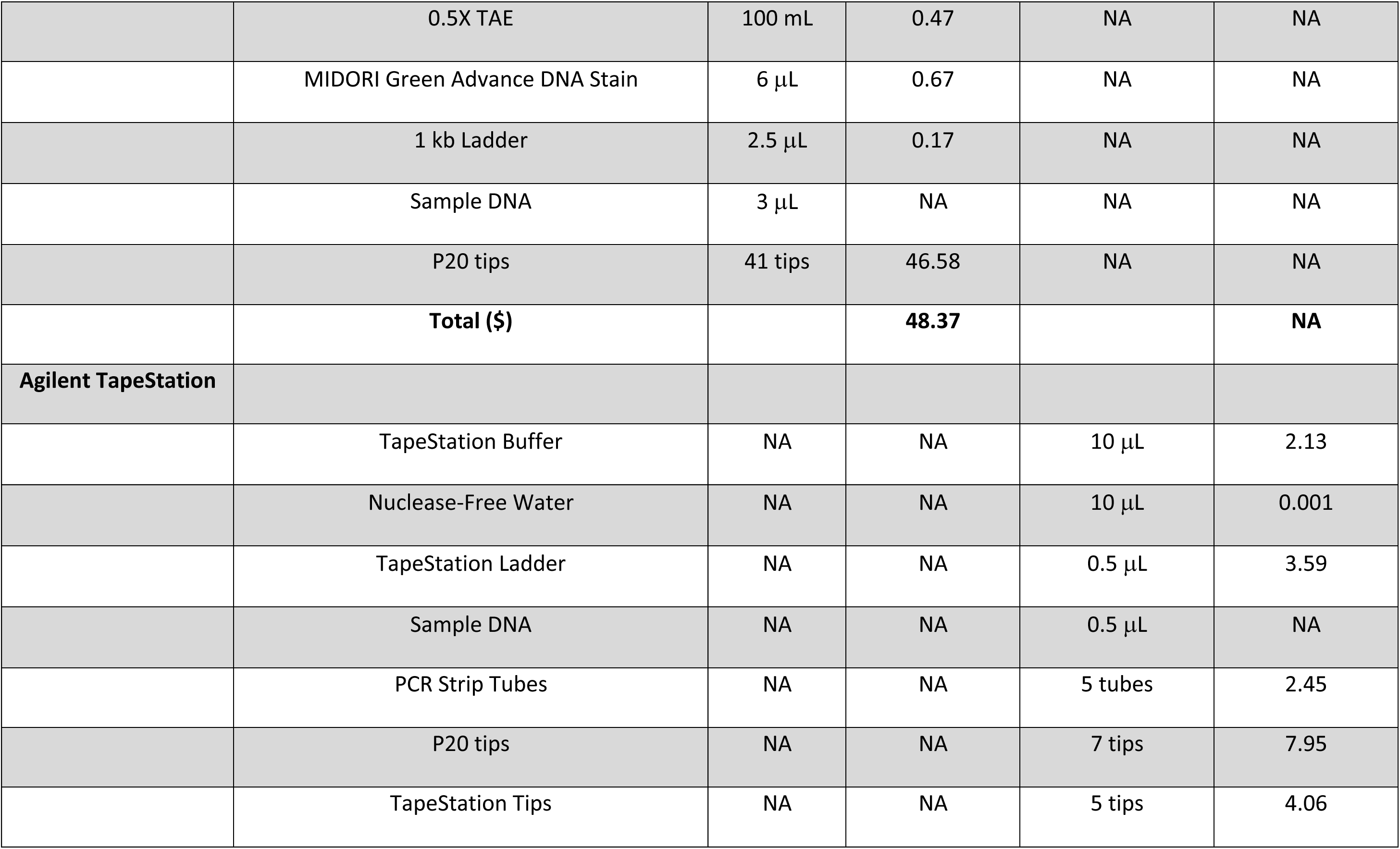
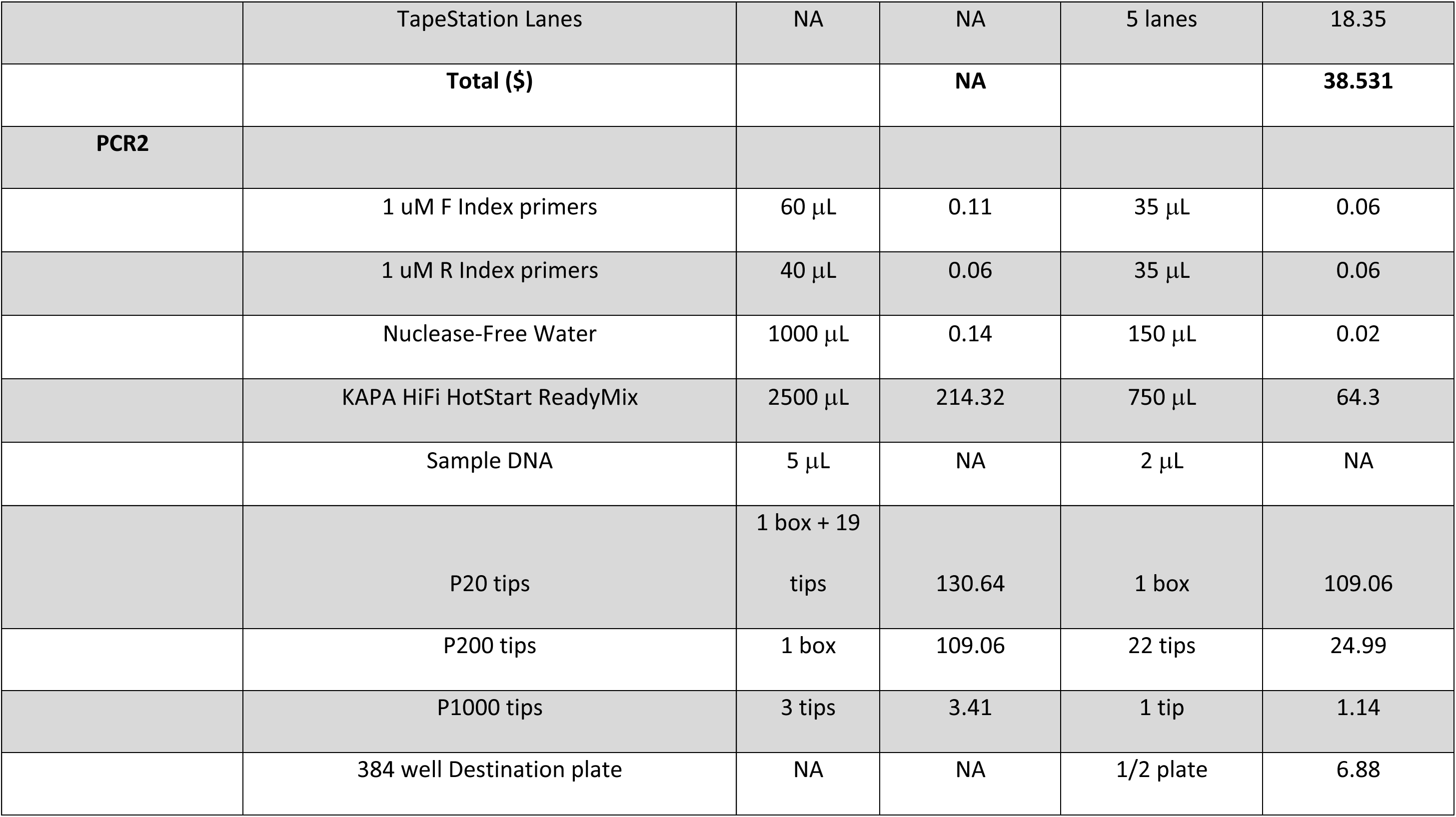
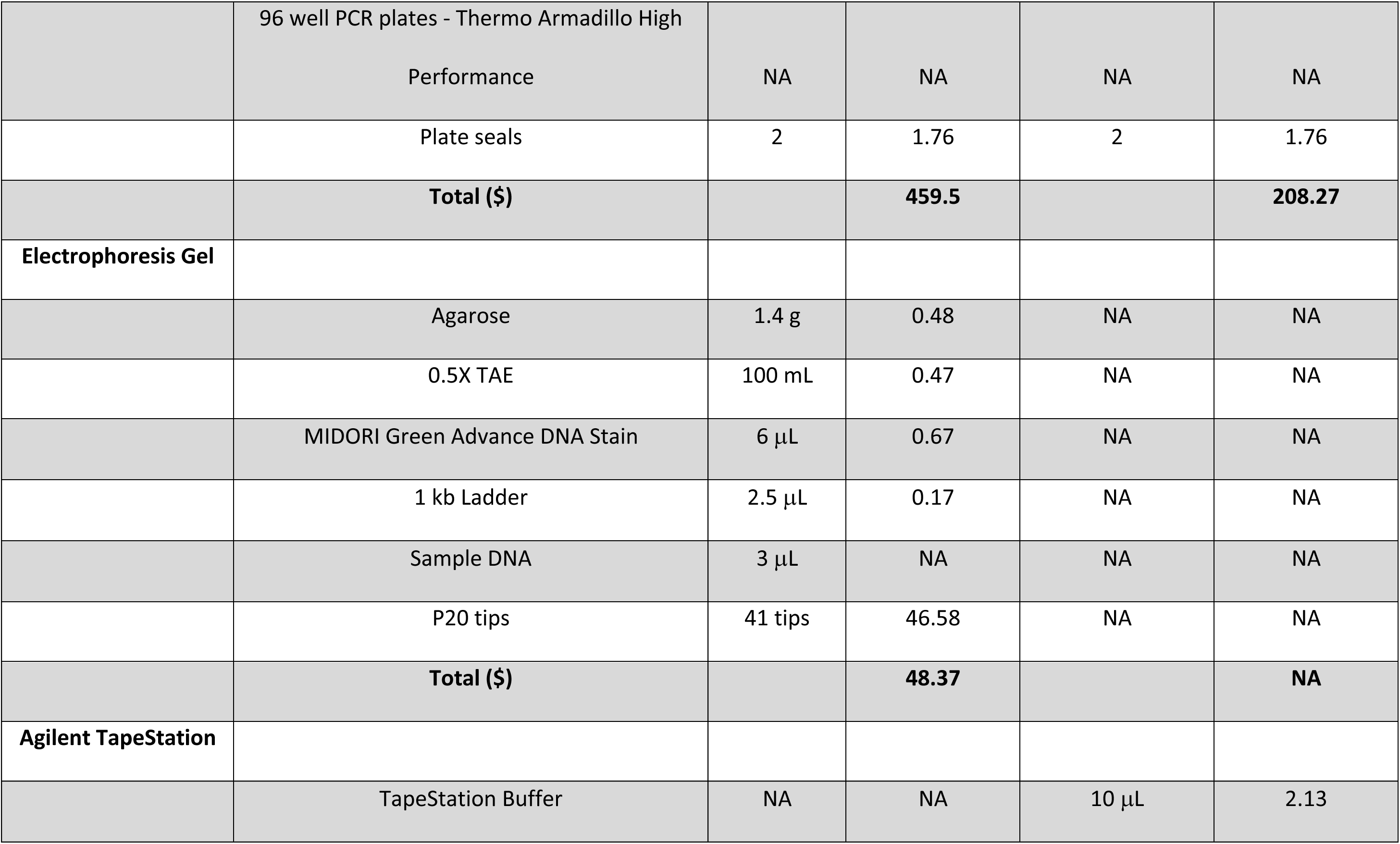
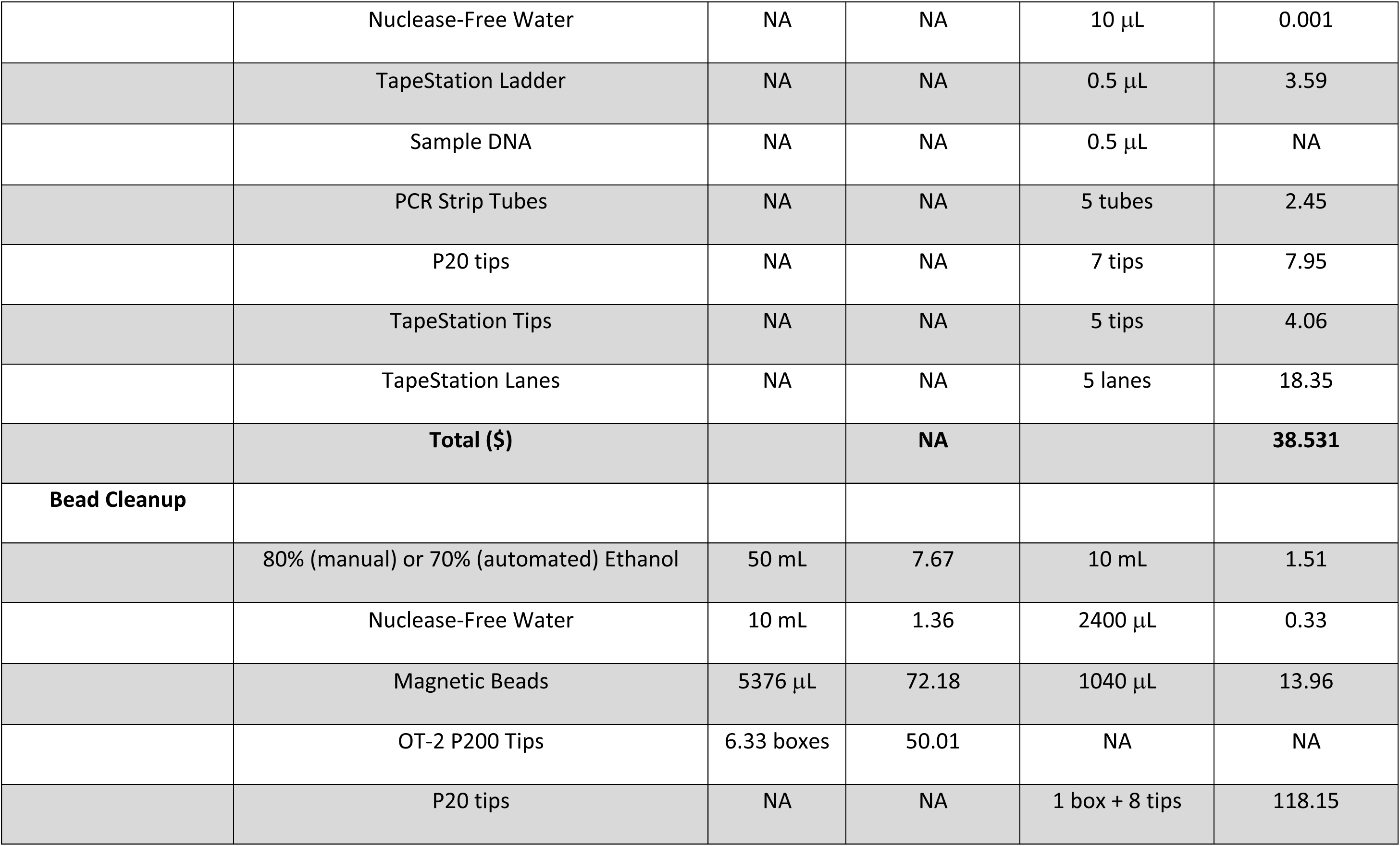
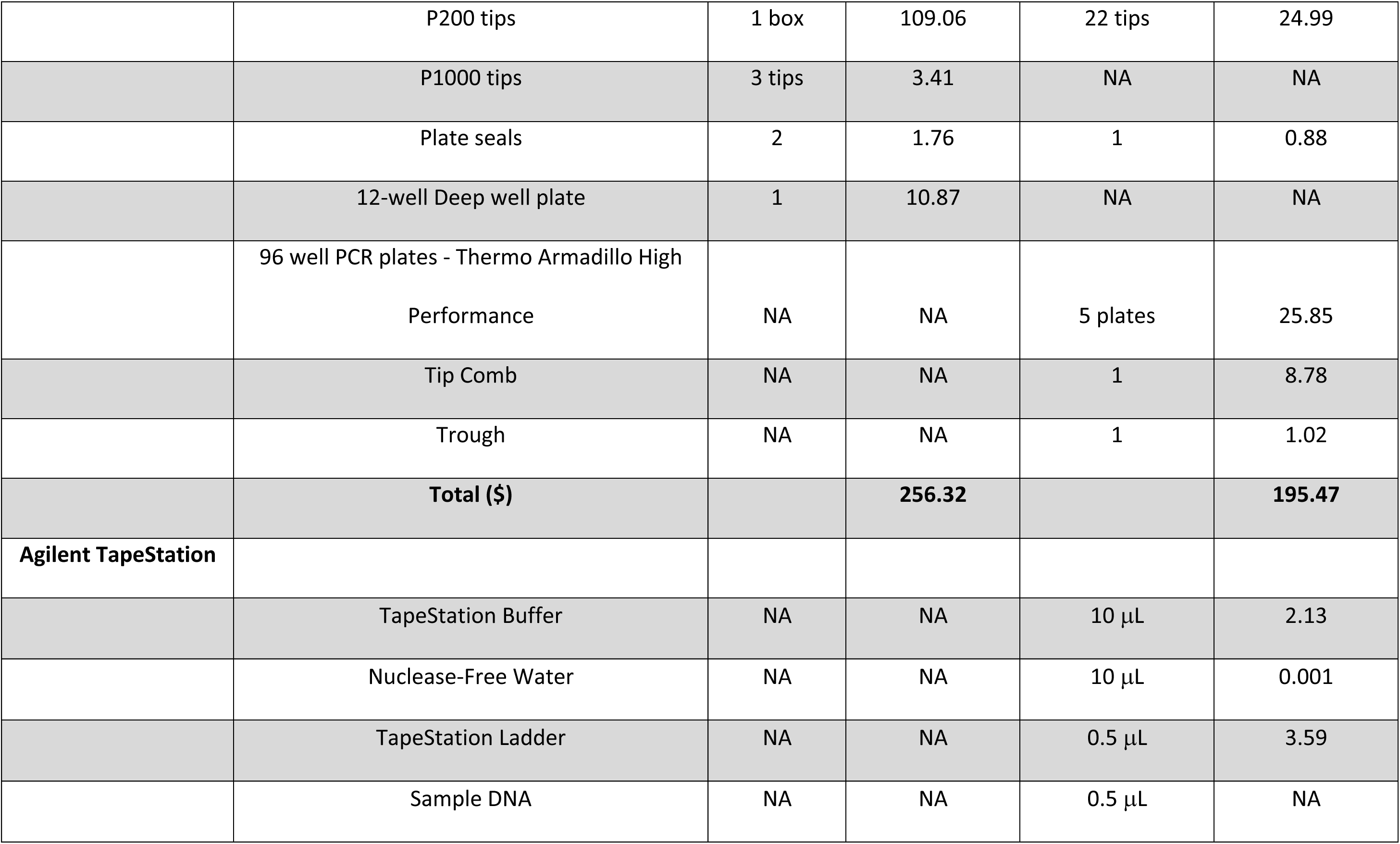
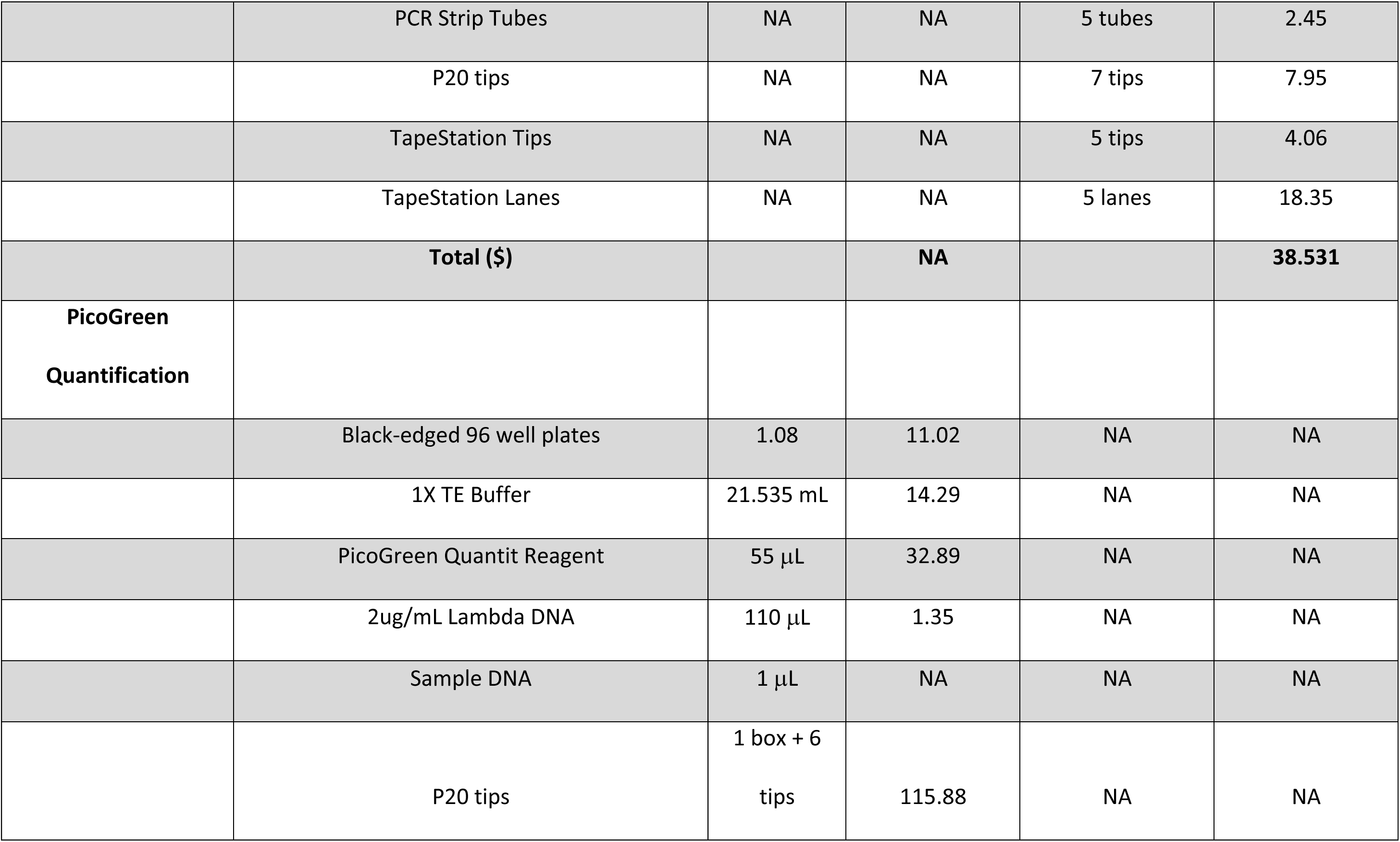
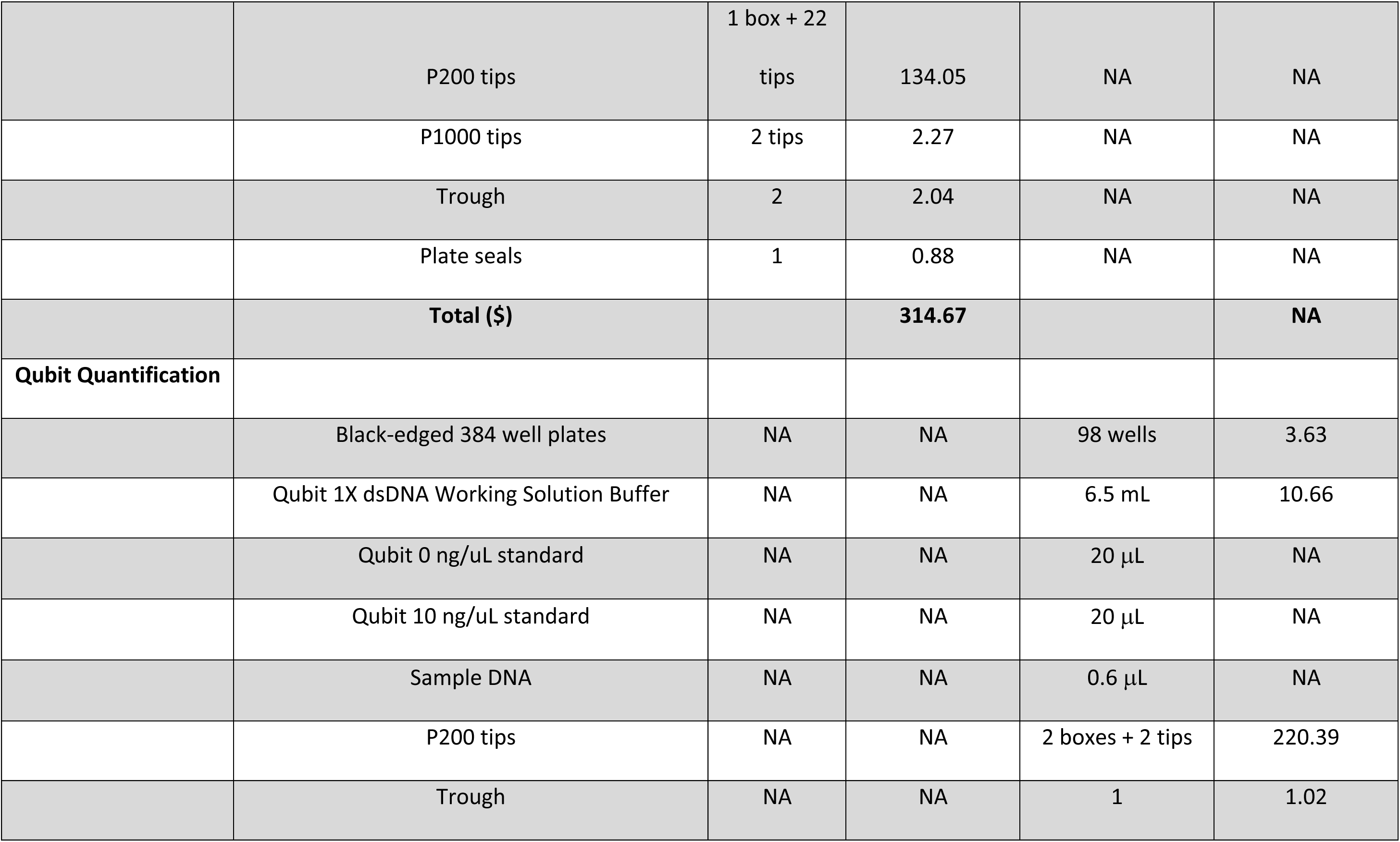
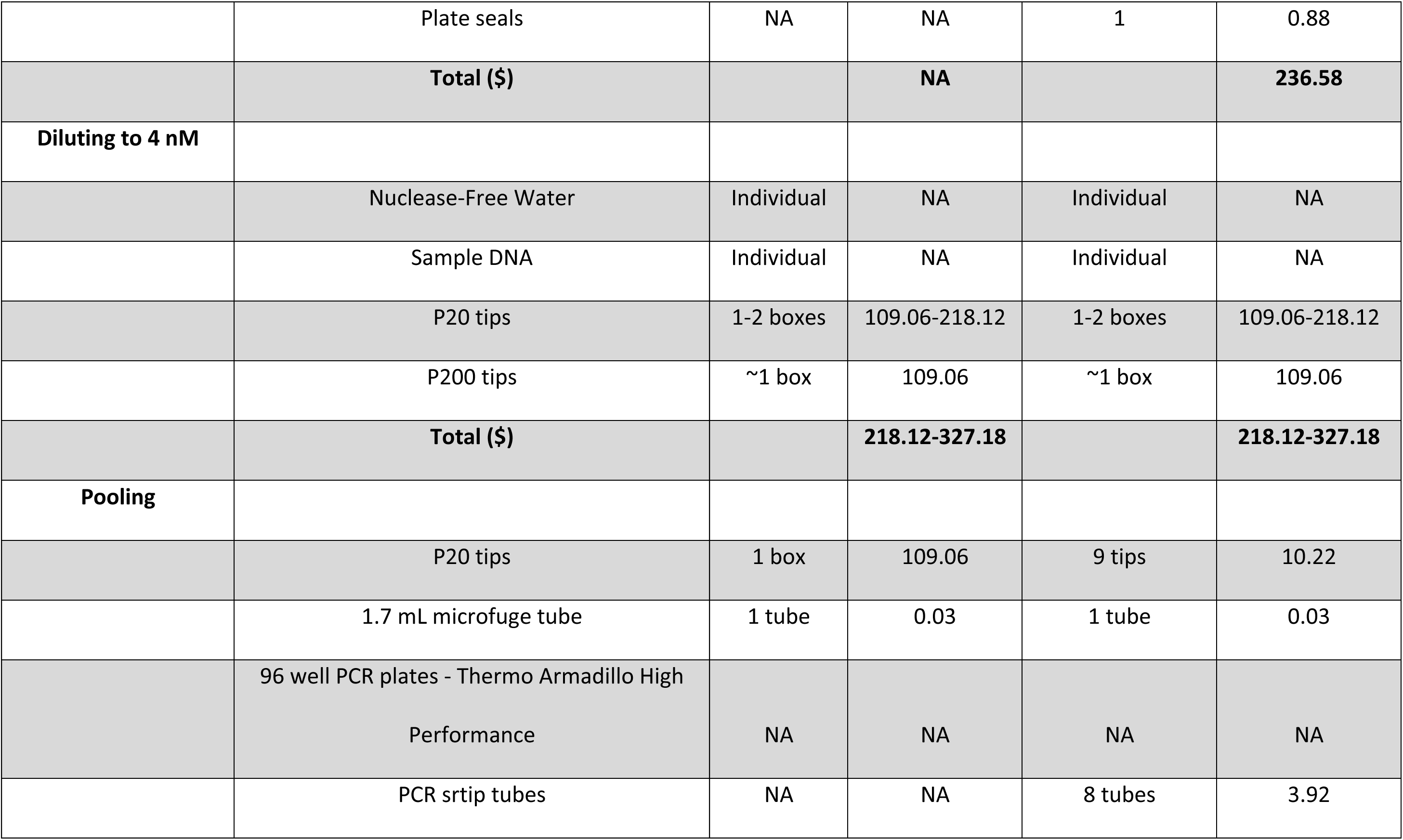
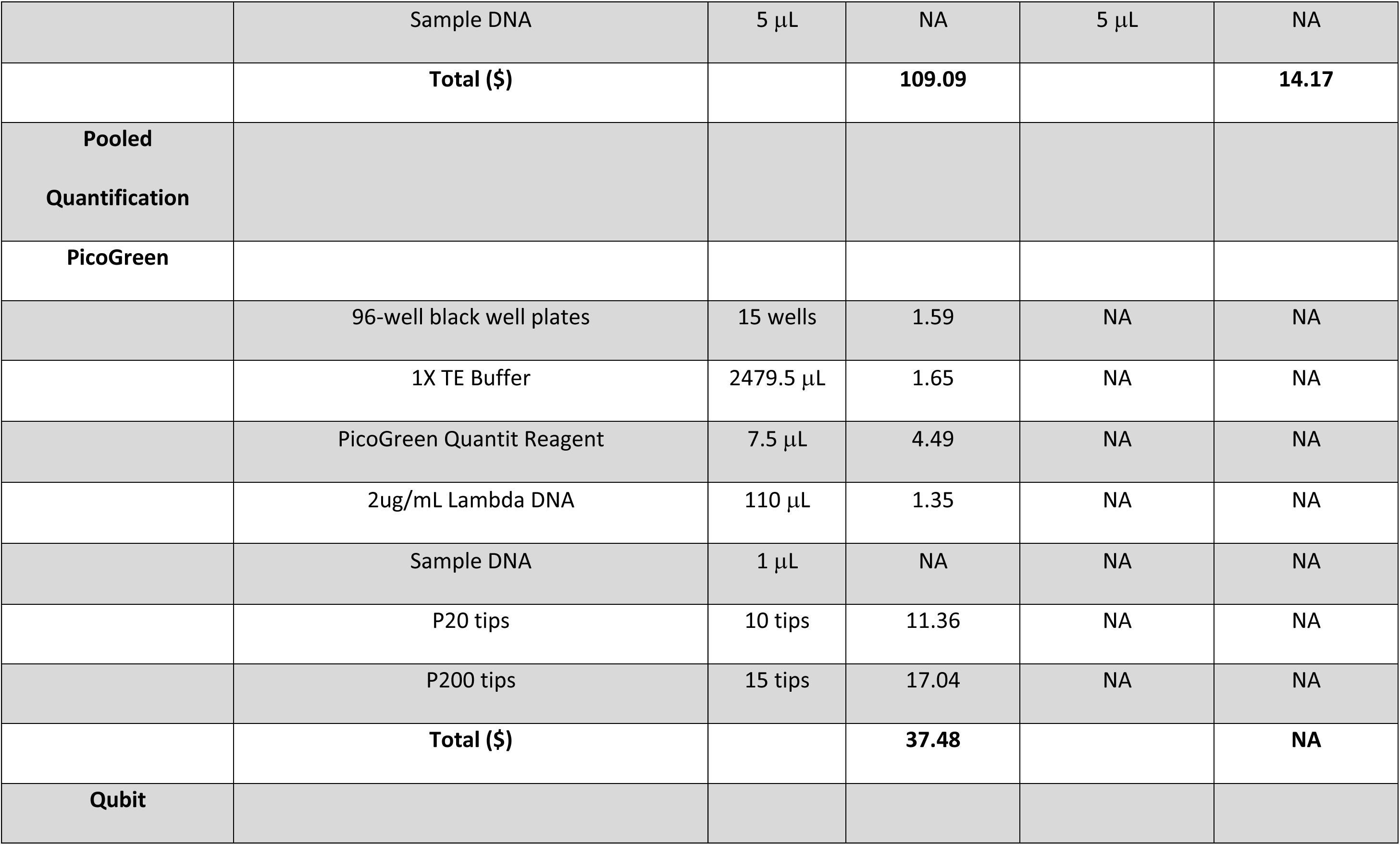
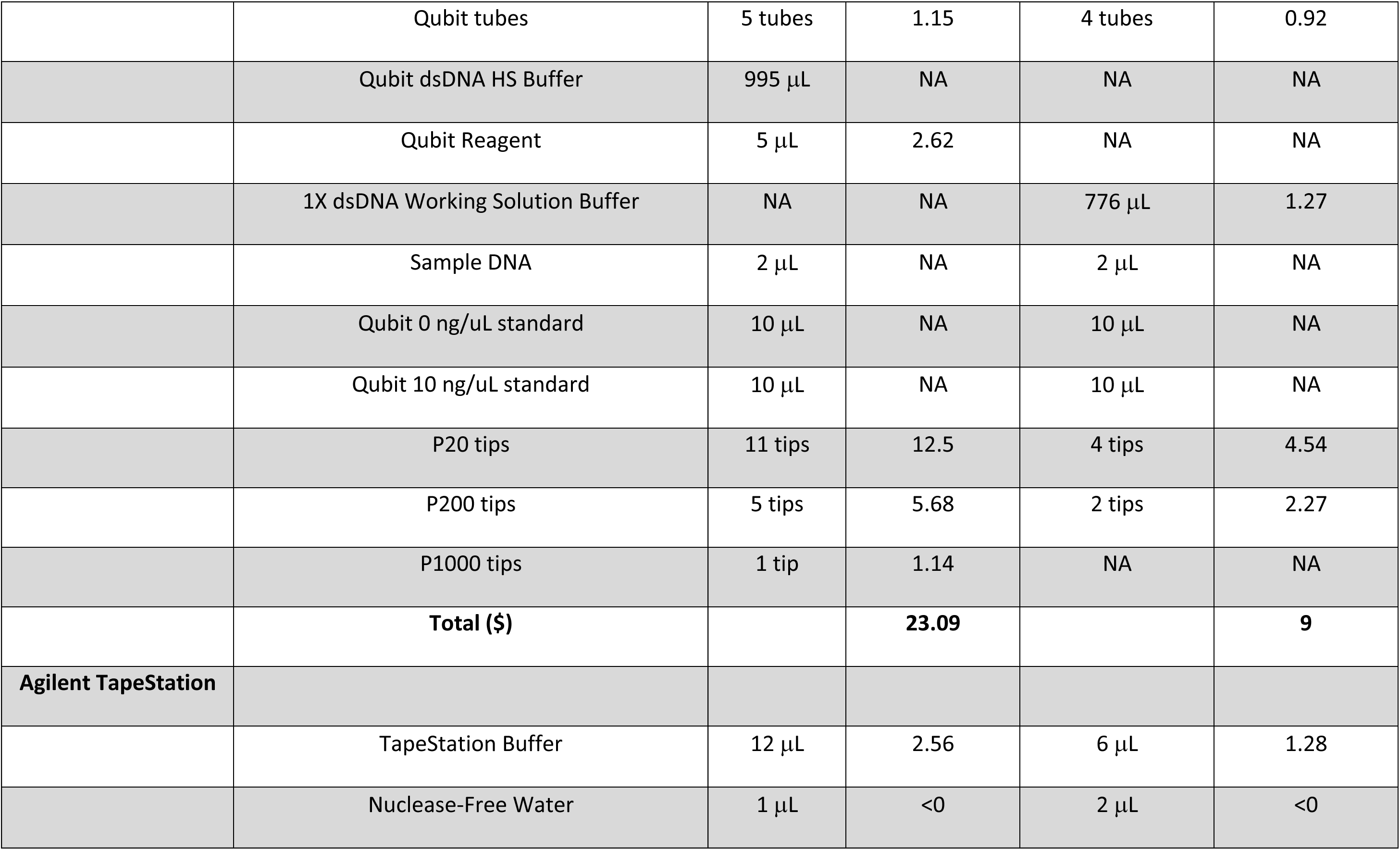
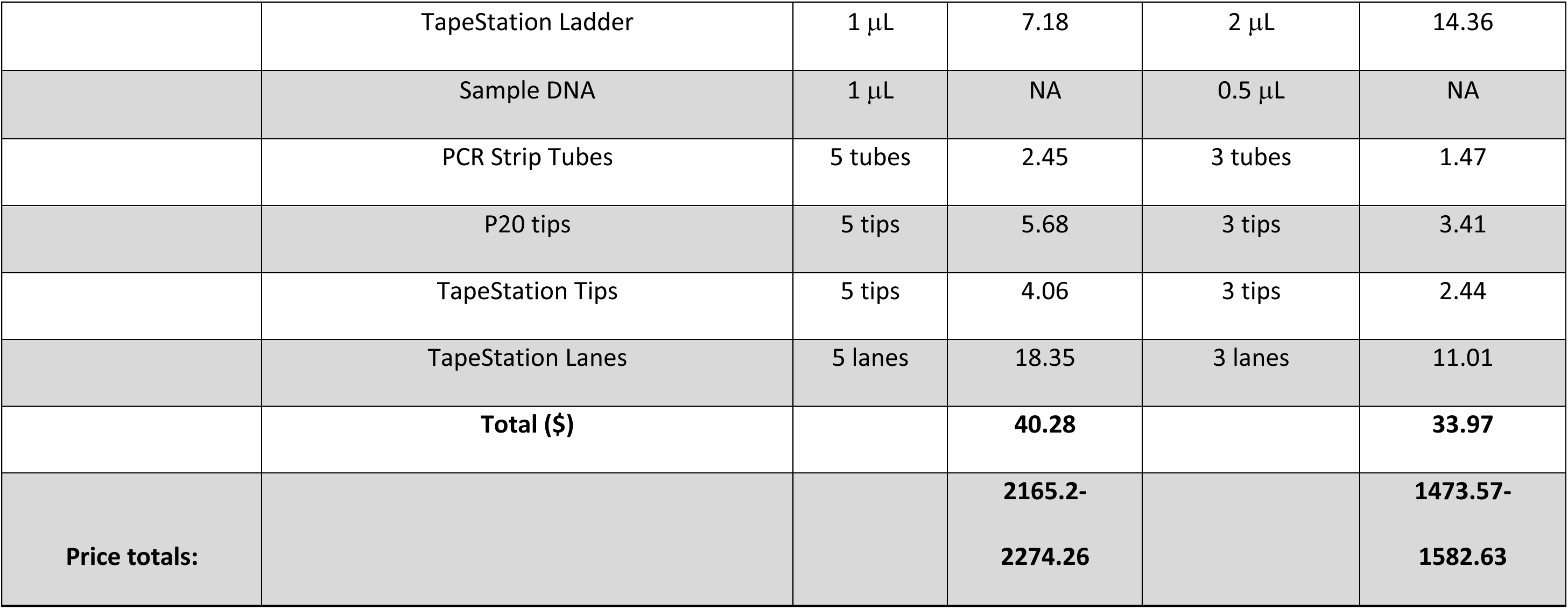
Table comparing consumables and reagents used for both manual and automated library preparation. for each step. The total price of each step and overall totals are given.

